# WHAMM functions in kidney reabsorption and polymerizes actin to promote autophagosomal membrane closure and cargo sequestration

**DOI:** 10.1101/2024.01.22.576497

**Authors:** Alyssa M Coulter, Valerie Cortés, Corey J Theodore, Rachel E Cianciolo, Ron Korstanje, Kenneth G Campellone

## Abstract

The actin cytoskeleton is essential for many functions of eukaryotic cells, but the factors that nucleate actin assembly are not well understood at the organismal level or in the context of disease. To explore the function of the actin nucleation factor WHAMM in mice, we examined how *Whamm* inactivation impacts kidney physiology and cellular proteostasis. We show that male WHAMM knockout mice excrete elevated levels of albumin, glucose, phosphate, and amino acids, and display abnormalities of the kidney proximal tubule, suggesting that WHAMM activity is important for nutrient reabsorption. In kidney tissue, the loss of WHAMM results in the accumulation of the lipidated autophagosomal membrane protein LC3, indicating an alteration in autophagy. In mouse fibroblasts and human proximal tubule cells, WHAMM and its binding partner the Arp2/3 complex control autophagic membrane closure and cargo receptor recruitment. These results reveal a role for WHAMM-mediated actin assembly in maintaining kidney function and promoting proper autophagosome membrane remodeling.

## INTRODUCTION

The actin cytoskeleton is crucial for controlling intracellular organization and the dynamics of membrane-bound organelles. To coordinate such cellular functions, globular (G-) actin monomers assemble into filamentous (F-) actin polymers (Pollard, 2016). Actin assembly is important in nearly all animal cells and tissues, although distinct physiological systems may rely on different regulatory factors (Rivers and Thrasher, 2017; Molinie and Gautreau, 2018; Kounakis and Tavernarakis, 2019). Despite this progress in understanding cytoskeletal activities, direct connections between dysfunctional actin assembly pathways and the pathogenesis of specific diseases are not well characterized.

To ensure that actin assembles when and where it is needed, proteins called nucleators direct the initiation of actin polymerization (Rottner *et al*., 2017; Gautreau *et al*., 2021). In mammals, these include a nucleator called the Arp2/3 complex, which cooperates with ∼12 activators, termed nucleation-promoting factors (Campellone and Welch, 2010). Most Arp2/3 activators are members of the Wiskott-Aldrich Syndrome Protein (WASP) family (Alekhina *et al*., 2017; Kabrawala *et al*., 2020). The WASP, WAVE, and WASH subgroups within the family have been thoroughly studied during plasma membrane dynamics, cell migration, and endocytic trafficking (Kramer *et al*., 2022). In contrast, the activities of the WHAMM/JMY subgroup have emerged in processes that were overlooked for many years (Campellone *et al*., 2023). WHAMM was discovered to promote ER-Golgi transport, endomembrane tubulation, and actin-microtubule interactions (Campellone *et al*., 2008; Shen *et al*., 2012; Russo *et al*., 2016), while JMY was recognized for roles in gene expression, motility, and *trans*-Golgi transport (Shikama *et al*., 1999; Zuchero *et al*., 2009; Schluter *et al*., 2014). More recent studies have revealed that WHAMM and JMY both function in autophagy and apoptosis (Coutts and La Thangue, 2015; Kast *et al*., 2015; Mathiowetz *et al*., 2017; Dai *et al*., 2019; Hu and Mullins, 2019; King *et al*., 2021; Wu *et al*., 2021). Although the activities of these two factors in apoptosis appear to lie in cytosolic actin rearrangements (King *et al*., 2021; King and Campellone, 2023), their participation in autophagy involves organelle remodeling.

Autophagy (formally, macroautophagy) is a mechanism of cytoplasmic digestion wherein double membrane-bound organelles called autophagosomes engulf cytoplasmic material and fuse with lysosomes for degradation (Zhao and Zhang, 2019; Vargas *et al*., 2023). This process is crucial for organismal development and cellular homeostasis and takes place constitutively, but is also induced by nutrient starvation, proteotoxic stress, and other stimuli (Dikic and Elazar, 2018; Levine and Kroemer, 2019). During autophagosome biogenesis, PI(3)P-rich phagophore membranes surround cytoplasmic cargo (Axe *et al*., 2008; Devereaux *et al*., 2013; Mi *et al*., 2015). This activity involves the ATG8 family of proteins, including the mammalian LC3s (LC3A/B/C) and GABARAPs (GABARAP, GABARAP-L1/L2), which exist as immature forms (e.g., LC3-I) that are cytosolic, and mature phosphatidylethanolamine-conjugated forms (e.g., LC3-II) that are physically linked to autophagosomal membranes (Mizushima, 2020; Klionsky *et al*., 2021). Selective autophagy receptors, such as SQSTM1/p62, act as adaptors by binding both to LC3 and ubiquitinated cellular ‘cargo’ (Pankiv *et al*., 2007; Johansen and Lamark, 2020; Vargas *et al*., 2023). Autophagic flux takes place upon syntaxin-mediated autophagosome fusion with lysosomes and the degradation and recycling of macromolecules (Nakamura and Yoshimori, 2017). Following autolysosomal membrane tubulation, lysosomes can be regenerated, enabling them to maintain cellular proteostasis (Ballabio and Bonifacino, 2020).

A specific function for actin dynamics in autophagy was initially revealed when the interiors of phagophores were found to be shaped by actin assembly in a PI(3)P-dependent manner (Mi *et al*., 2015). Subsequently, WHAMM and JMY were shown to act at multiple steps in the canonical autophagy pathway. WHAMM binds to PI(3)P, localizes to subdomains of nascent autophagosomes, and is important for efficient LC3 lipidation (Mathiowetz *et al*., 2017). WHAMM-driven Arp2/3 activation also increases the size of autophagosomes and causes actin-based rocketing in the cytosol (Kast *et al*., 2015). JMY binds LC3 and affects autophagosome maturation (Coutts and La Thangue, 2015). Upon activation by LC3, JMY additionally promotes actin-based autophagosome rocketing (Hu and Mullins, 2019). WHAMM participates again later in the autophagy pathway by binding PI(4,5)P2 and mediating autolysosome tubulation (Dai *et al*., 2019; Wu *et al*., 2021). WHAMM function is ultimately important for the degradation of p62 and turnover of ubiquitinated cargo (Mathiowetz *et al*., 2017).

Although the founding member of the WASP family was discovered due to genetic mutations in patients with immunodeficiencies decades ago (Derry *et al*., 1994), surprisingly little is understood about how alterations in other family members contribute to disease. Several mutant versions of WAVE- or WASH-binding proteins have been observed in individuals with neurological or immunological disorders (Kramer *et al*., 2022; Campellone *et al*., 2023), but mutations in the WASP-family genes themselves are only beginning to be characterized (Valdmanis *et al*., 2007; Ropers *et al*., 2011; Ito *et al*., 2018; Courtland *et al*., 2021; Srivastava *et al*., 2021). *WHAMM* variants play a potential role in disease, as most Amish patients with the rare neurodevelopmental/kidney disorder Galloway-Mowat Syndrome (GMS) harbor a homozygous *WHAMM* mutation that abrogates WHAMM-driven Arp2/3 activation *in vitro* and leads to autophagy defects in cells (Jinks *et al*., 2015; Mathiowetz *et al*., 2017). However, GMS has a complex genetic basis, as Amish patients also possess a homozygous mutation in the nearby *WDR73* gene which is considered to be disease-causing (Jinks *et al*., 2015). *WDR73* mutations are associated with multiple neurological illnesses (Colin *et al*., 2014; Ben-Omran *et al*., 2015; Vodopiutz *et al*., 2015; Jiang *et al*., 2017; El Younsi *et al*., 2019; Tilley *et al*., 2021), and loss-of-function mutations in many different genes can give rise to GMS or GMS-like conditions (Braun *et al*., 2017; Rosti *et al*., 2017; Braun *et al*., 2018; Arrondel *et al*., 2019; Mann *et al*., 2021). Thus, the contribution of WHAMM to health and disease is difficult to discern. In the current study, we generated a null mutation in mouse *Whamm* to better define its role in kidney physiology and cellular autophagy.

## RESULTS

### WHAMM knockout male mice display proximal tubule reabsorption defects

To understand the organismal function of WHAMM, we used a *Whamm* allele with a targeted deletion in exon 3 to generate homozygous WHAMM knockout (WHAMM^KO^) mice (Figure 1A). We then compared the mutant mice to wild type (WHAMM^WT^) littermates in multiple phenotypic analyses. Since Amish GMS patients have kidney abnormalities resulting in proteinuria (Jinks *et al*., 2015), we tested the urine from wild type and knockout mice for albuminuria. WHAMM^KO^ males displayed a significant increase in the albumin-to-creatinine ratio (ACR) compared to WHAMM^WT^ males at 24 weeks-of-age (Figure 1B). In contrast, ACRs for KO and WT females were statistically indistinguishable from one another (Figure 1B). To determine the age of onset for male albuminuria, we compared the ACRs at 16, 20, and 24-week timepoints. While WHAMM^WT^ males showed a slight decrease in ACR over time, WHAMM^KO^ males showed a gradual increase in ACR, although a statistically significant difference between the WT and KO was not reached until 24 weeks (Supplemental Figure S1).

**Figure 1.**
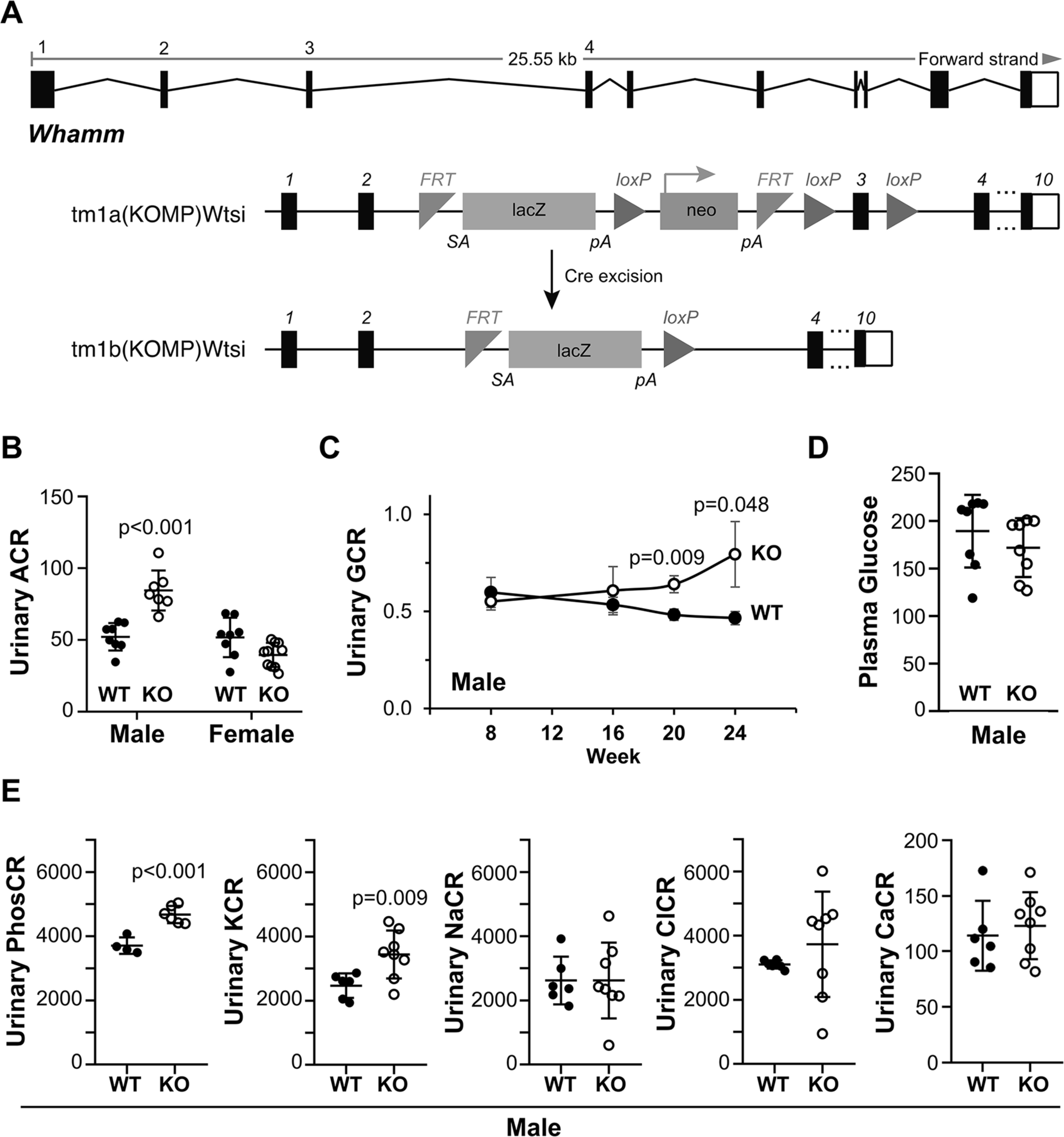
Male WHAMM^KO^ mice excrete elevated levels of albumin, glucose, phosphate, and potassium in their urine. **(A)** The mouse *Whamm* gene is 25kb in length and contains 10 exons. A floxed allele containing IRES-*lacZ* and neo cassettes between exons 2 and 3, as well as *loxP* sites between the cassettes and flanking exon 3 was generated. Following Cre-mediated recombination, a knockout allele was created. Internal ribosome entry site (IRES); splice acceptor (SA); polyadenylation (pA); flippase recombination target (FRT). **(B)** Urine samples were collected from male and female wild type (WT; filled circles) or WHAMM knockout (KO; open circles) mice at 24 weeks-of-age and subjected to urinalysis. Urinary albumin-to-creatinine (ACR) ratios are plotted. Each circle represents one mouse. Statistical bars display the mean ±SD from n=7-10 mice. **(C)** Urinary glucose-to-creatinine (GCR) ratios for males from 8 to 24 weeks-of-age are plotted. Each circle represents the mean ±SE from n=6-9 mice per genotype for each timepoint. **(D)** Plasma glucose-to-creatinine ratios for males at 24 weeks-of-age are plotted. Statistical bars display the mean ±SD from n=8 mice. **(E)** Urinary phosphate (PhosCR), potassium (KCR), sodium (NaCR), chloride (ClCR), and calcium (CaCR) to creatinine ratios are plotted. Each circle represents one male mouse at 24 weeks. Statistical bars display the mean ±SD from n=4-8 mice. Significant p-values are noted (unpaired t-tests).

To explore whether the inactivation of *Whamm* affected other parameters of kidney function, we next analyzed urinary glucose levels. WHAMM^KO^ male mice displayed significantly higher glucose-to-creatinine ratios (GCR) at 20 and 24 weeks-of-age (Figure 1C). This phenotype was also sex-specific, as knockout and wild type females were similar to one another (Supplemental Figure S1). The urinary excretion of glucose in males did not appear to be caused by diabetes, as non-fasting plasma glucose levels did not differ between the KO and WT (Figure 1D).

Given the potential loss of multiple molecules in urine, we additionally measured urinary phosphate, potassium, sodium, chloride, and calcium in the 24-week-old males. The WHAMM^KO^ mice had a significantly higher urinary phosphate-to-creatinine ratio (PhosCR) and potassium-to-creatinine ratio (KCR) than the WHAMM^WT^ mice, whereas urinary sodium (NaCR), chloride (ClCR), and calcium (CaCR) did not show differences between the two genotypes of male mice (Figure 1E). Because glucose, phosphate, and potassium are reabsorbed from the filtrate in the proximal tubule, these results are indicative of a tubular malfunction in the WHAMM^KO^ males.

Notably, the WHAMM^KO^ excretion phenotypes are reminiscent of those found in renal Fanconi Syndromes, proximal tubule diseases with diverse genetic bases (Klootwijk *et al*., 2015; Lemaire, 2021). As amino aciduria is also seen in Fanconi Syndromes, we used mass spectrometry to measure the relative levels of different amino acids in the urine of 24-week-old knockout and wild type males. The WHAMM^KO^ males showed statistically significant increases in urinary glutamic acid, glutamine, methionine, pipecolic acid, proline, and cysteine (Figure 2). These results further support the conclusion that WHAMM deficiency causes a Fanconi-like Syndrome in male mice.

**Figure 2.**
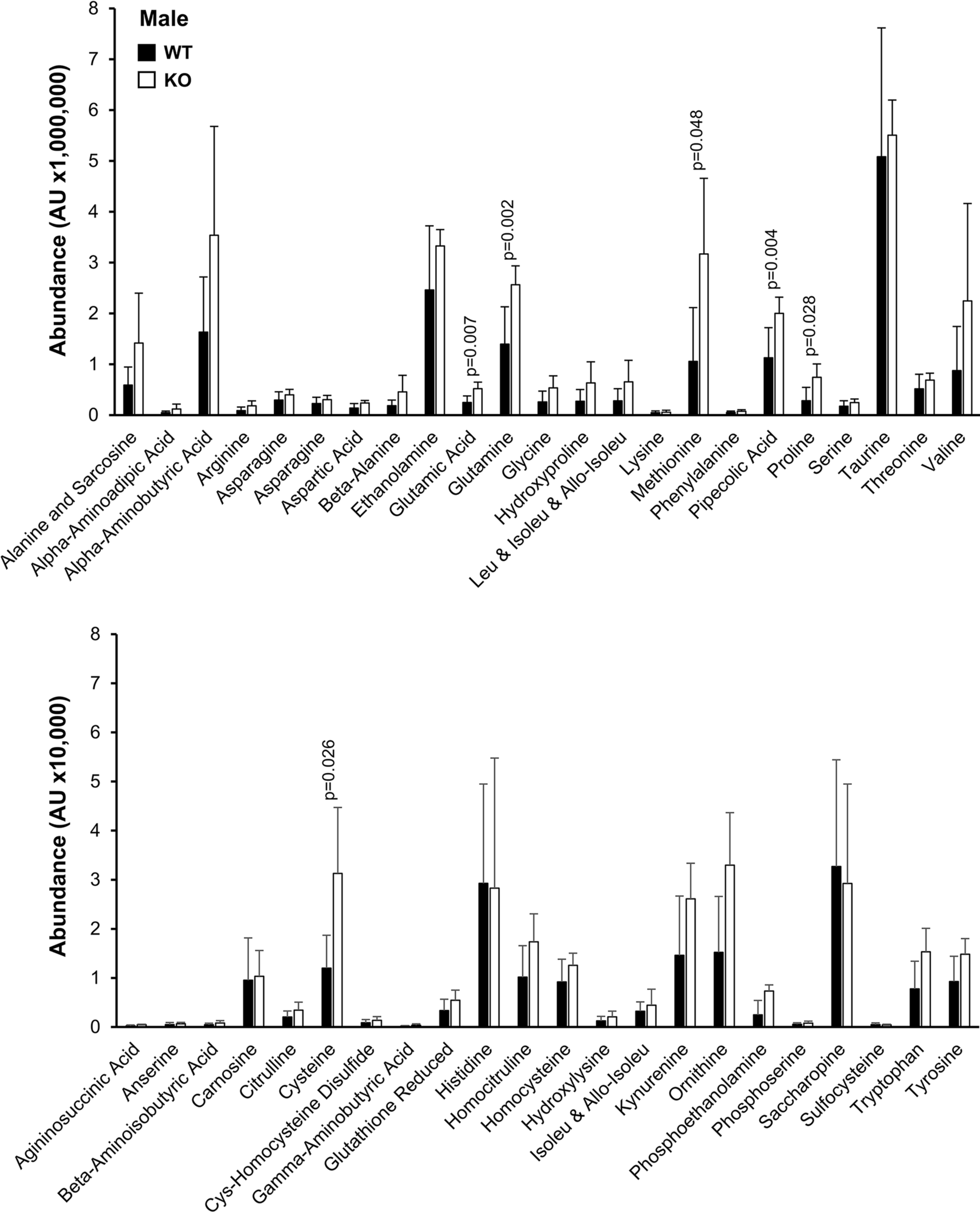
Male WHAMM^KO^ mice excrete elevated levels of amino acids in their urine. Urine samples were collected from male wild type (WT; black bars) or WHAMM knockout (KO; white bars) mice at 24 weeks-of-age, and urinary amino acids were measured using mass spectrometry. AU = arbitrary units. Each bar represents the mean ±SD from n=5-6 mice. Significant p-values are noted (ANOVA).

### WHAMM deletion alters proximal tubule polarity in male kidneys

Previous work has shown that WHAMM protein is abundant in the human and mouse kidney (Campellone *et al*., 2008), but the more precise locations of its expression in the nephron have not been described. To assess the cell type-specific mRNA expression pattern of *Whamm*, we surveyed single-cell sequencing data from male and female adult mouse kidneys (Ransick *et al*., 2019). *Whamm* mRNA was present throughout the proximal tubule of both male and female mice, with its highest expression found in segment 2 of the proximal tubule of male mice (Supplemental Figure S2). *Whamm* levels in the proximal tubule were higher than those in podocytes but less than those in intercalated type-B cells of the cortical collecting duct (Supplemental Figure S2). Analyses of *Whamm* and other WASP-family and Arp2/3 complex genes indicated that they were all expressed in podocytes and proximal tubules to varying degrees, with the exceptions of *Was* (encoding WASP), *Wasf1* (WAVE1), and *Actr3b* (the Arp3B isoform), which were absent (Supplemental Figure S3). WHAMM expression in the proximal tubule of male mice is therefore amenable to a role in tubular reabsorption.

Some proximal tubule disorders have been attributed to endocytic trafficking defects and reductions in the quantities of receptor proteins (Norden *et al*., 2002; Oltrabella *et al*., 2015; Inoue *et al*., 2017; Festa *et al*., 2019; Berquez *et al*., 2020; Lemaire, 2021). Given that the physiological abnormalities in WHAMM-deficient mice were sex-specific, we focused our efforts on characterizing tissues and cells from males. To assess the amounts of the multiligand endocytic receptor LRP2/Megalin, the tubule-expressed angiotensin-converting enzyme ACE2, the glucose transporter SGLT2, and the phosphate transporter SLC20A1 in WHAMM^WT^ and WHAMM^KO^ males, we isolated their kidneys and generated tissue extracts for immunoblotting. While Megalin levels were highly variable, especially in the knockouts, the relative amount of each receptor protein was generally similar in WHAMM^WT^ and WHAMM^KO^ kidney tissue (Supplemental Figure S4), suggesting that WHAMM deficiency does not dramatically alter the abundance of membrane receptors in the kidney.

To provide a broad appraisal of kidney structure in the mice, we performed histological analyses of kidney sections after periodic acid-Schiff (PAS) staining, but did not observe any major differences between samples from male WHAMM^WT^ and WHAMM^KO^ animals (all slides can be viewed at https://images.jax.org/webclient/?show=dataset-2763). To more specifically visualize proximal tubule morphology and polarity, we stained kidney sections from WHAMM^WT^ and WHAMM^KO^ males with fluorescent antibodies to Megalin and ACE2 (Figure 3A; Supplemental Figure S5), as both proteins are expected to localize to the apical regions of proximal tubule cells (Kerjaschki *et al*., 1984; Warner *et al*., 2005).

**Figure 3.**
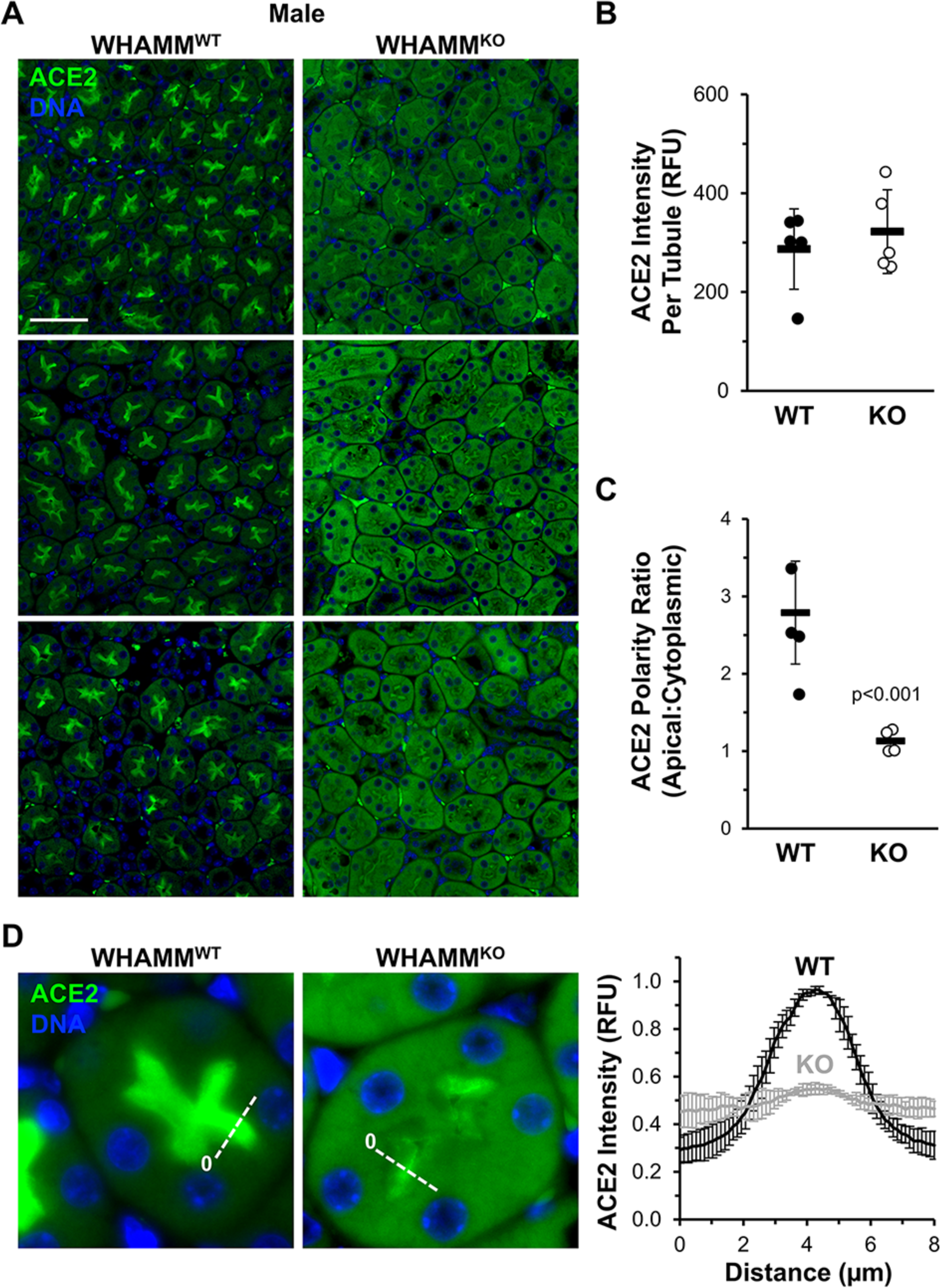
Polarized ACE2 staining in the kidney proximal tubule is reduced in male WHAMM^KO^ mice. **(A)** Kidney tissue sections from wild type (WHAMM^WT^) or WHAMM knockout (WHAMM^KO^) male mice were stained with ACE2 antibodies (green) and DAPI (DNA; blue). Scale bar, 50µm. **(B)** The ACE2 fluorescence intensity per tubule was calculated in ImageJ. Each circle represents the average ACE2 kidney staining from an individual mouse in which approximately 75 tubules were examined. Statistical bars represent the mean ±SD from n=5 mice. **(C)** The ACE2 polarity ratio was calculated in ImageJ by dividing the fluorescence intensity in an apical region of the tubule by the intensity in a cytoplasmic region. Each circle represents the average ratio from an individual mouse in which 40 tubules were examined. Statistical bars represent the mean ±SD from n=4 mice. **(D)** Kidney tissue sections from (A) were used to generate pixel intensity profiles. Lines were drawn through the center of the tubule, and the ACE2 intensity along the line was plotted. The origin of each line is indicated with a 0. Plotted points represent the normalized mean ACE2 fluorescence ±SD from n=3 mice per genotype (comprising 3 pixel intensity plots per mouse). RFU = relative fluorescence units. Significant p-values are noted (unpaired t-tests).

In accordance with the earlier immunoblotting results, measurements of the immunofluorescence intensities of Megalin and ACE2 revealed that wild type and knockout males contained similar amounts of each protein per cluster of tubular cells (Figure 3B; Supplemental Figure S5). However, close inspection of the localization of each protein revealed several differences. Megalin immunostaining was quite variable, but exhibited a primarily apical localization in most WHAMM^WT^ samples (Supplemental Figure S5). In WHAMM^KO^ samples, tubular clusters looked more disorganized and sometimes displayed a Megalin staining pattern that was less apical and more cytoplasmic (Supplemental Figure S5). For ACE2, staining was consistently apical in WT kidney sections but strikingly cytoplasmic in the KO sections (Figure 3A). Quantification of apical and cytoplasmic ACE2 intensity across many tubule clusters in multiple animals demonstrated that ACE2 displayed an apical-to-cytoplasmic polarity ratio of 3:1 in WHAMM^WT^ samples but that the polarity ratio fell to nearly 1:1 for the WHAMM^KO^ (Figure 3C). A defect in polarized receptor distribution was further confirmed using fluorescence linescan analyses, which showed sharp peaks of apical ACE2 intensity in WHAMM^WT^ kidneys but a muted apical ACE2 localization WHAMM^KO^ kidneys (Figure 3D). Together, our kidney tissue immunoblotting and immunofluorescence data demonstrate that while receptor abundance is relatively normal across both genotypes of male mice, the organization within proximal tubule cells appears to be distorted in the absence of WHAMM.

### The lipidation status of the autophagosomal protein LC3 is altered in WHAMM^KO^ kidneys

Cellular proteostasis systems are important for maintaining the integrity of proximal tubules (Cybulsky, 2017; Tang *et al*., 2020), and conditional deletions of Vps34, a key initiator of autophagic membrane biogenesis, alter the urinary proteome and perturb the apical localization of several membrane transport proteins including ACE2 in mice (Grieco *et al*., 2018; Rinschen *et al*., 2022). Because WHAMM plays a role in multiple steps of autophagy (Kast *et al*., 2015; Mathiowetz *et al*., 2017; Dai *et al*., 2019; Wu *et al*., 2021), we next sought to determine whether some aspect of autophagy might be altered in WHAMM^KO^ kidney tissue. The LC3 and GABARAP families of proteins are the most widely accepted markers of autophagosomal membranes, and LC3-II and GABARAP-II levels are considered to correlate with autophagosome quantities (Klionsky *et al*., 2021), so we immunoblotted kidney extracts with polyclonal antibodies that recognize both the immature and mature species of multiple LC3 and GABARAP isoforms.

In male kidneys, immature LC3-I and GABARAP-I levels appeared similar across both WHAMM genotypes (Figure 4A). However, mature lipidated LC3-II was more abundant in WHAMM^KO^ than WHAMM^WT^ kidney tissue (Figure 4A). None of the kidney extracts contained detectable levels of GABARAP-II (Figure 4A). Quantification of the LC3 species revealed that LC3-II was present, on average, in 3-fold higher amounts in the knockout than the wild type, and that the LC3 II:I ratio was also substantially greater in the KO males (Figure 4B). In female kidneys, LC3-II levels were not elevated. The only statistically significant difference between KO and WT females was in the LC3 II:I ratio, which was slightly reduced in the WHAMM^KO^ (Supplemental Figure S6). Collectively, these results suggest that the lack of WHAMM leads to altered LC3 modification in the murine kidney, with males experiencing an increase in LC3 lipidation and/or a decrease in LC3-II turnover.

**Figure 4.**
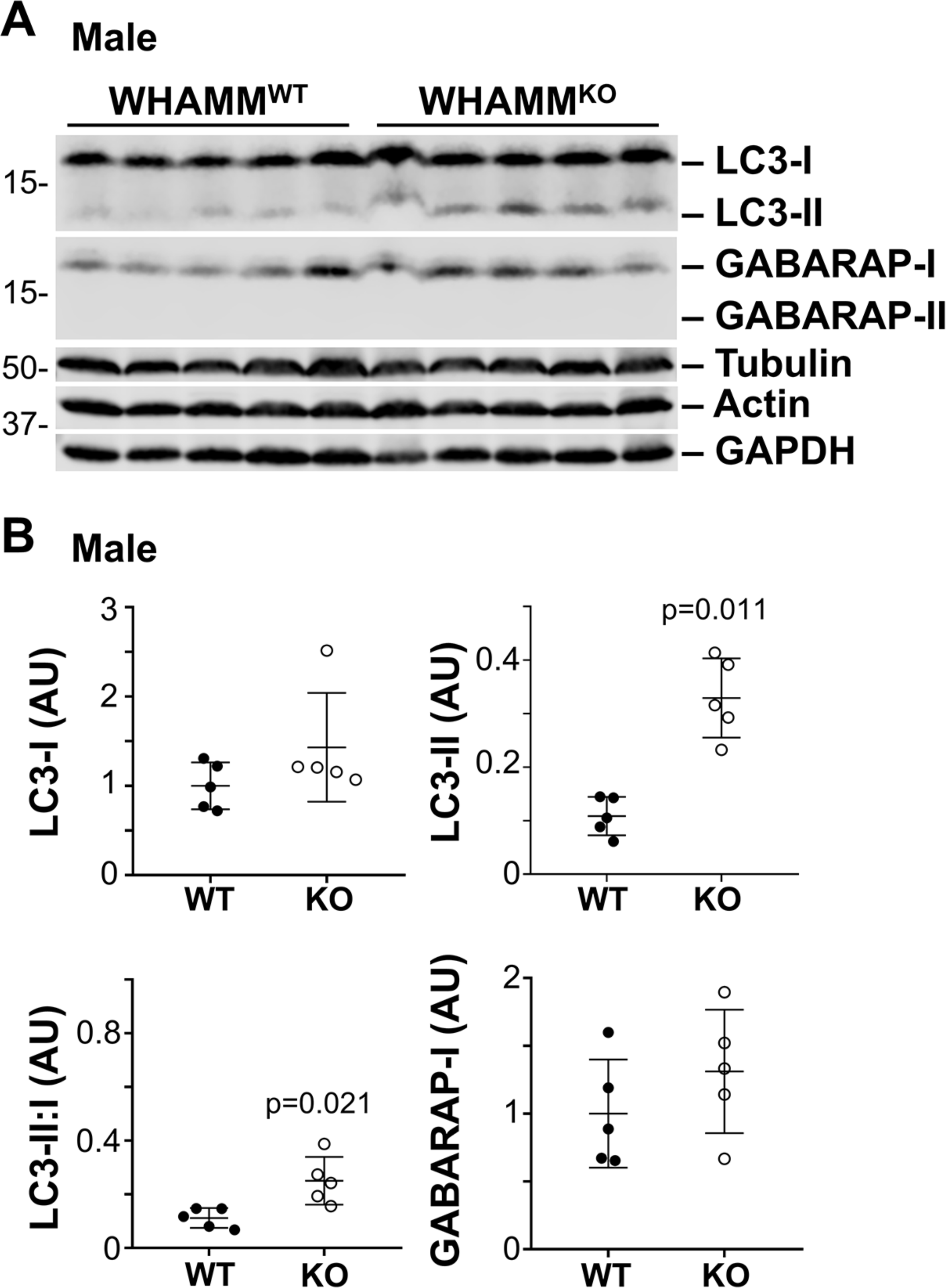
The lipidated form of the autophagosomal protein LC3 is more abundant in male WHAMM^KO^ kidneys. **(A)** Kidneys were harvested from 5 male WHAMM^WT^ and 5 male WHAMM^KO^ mice. 50µg extract samples were subjected to SDS-PAGE and immunoblotted with antibodies to LC3, GABARAP, tubulin, actin, and GAPDH. **(B)** LC3 and GABARAP band intensities in (A) were quantified relative to tubulin, actin, and GAPDH, and the mean normalized values were plotted. The LC3-II:I ratio was calculated by dividing the LC3-II band intensity by the LC3-I band intensity within each non-normalized sample. Statistical bars represent the mean ±SD from n=5 mice. Significant p-values are noted (unpaired t-tests).

### The morphogenesis of autophagic membranes is controlled by WHAMM

To better define the function of WHAMM at the cellular level, we next generated male WHAMM- proficient (WHAMM^HET^) and WHAMM-deficient (WHAMM^KO^) mouse embryonic fibroblasts (MEFs) (Supplemental Figure S7). WHAMM was initially characterized for its ability to activate Arp2/3 complex-dependent actin assembly, bind microtubules, and interact with membranes to promote anterograde transport (Campellone *et al*., 2008), but F-actin, microtubule, and *cis*-Golgi organization was relatively normal in knockout MEFs under standard culture conditions (Supplemental Figure S7).

Given our findings that WHAMM-deficient mouse kidneys accumulated lipidated LC3, we examined how WHAMM deletion affected autophagy in MEFs. At steady state, MEFs displayed diffuse LC3 and GABARAP staining without any cytosolic puncta and with minimal differences between genotypes (Supplemental Figure S7). This could be the result of low basal levels of autophagy and/or high rates of autophagosome turnover in embryo-derived cells. Therefore, to visualize autophagic structures, we prevented lysosomal degradation by treating MEFs with chloroquine. While WHAMM^HET^ cells formed discrete cup-shaped and ring-like LC3- and GABARAP-positive structures reminiscent of autophagosomes, WHAMM^KO^ cells showed more diffuse and small punctate LC3 and GABARAP staining patterns (Figure 5, A and B). Quantification of whole cell mean fluorescence intensities for LC3 and GABARAP demonstrated that the WHAMM deletion increased the intracellular abundance of both ATG8 subfamilies (Figure 5C). In addition, whereas actin localized to LC3- and GABARAP-labeled membranes in WHAMM^HET^ MEFs, little actin was recruited to autophagosomes in WHAMM^KO^ MEFs (Figure 5, A and B). These findings indicate that the permanent loss of WHAMM in MEFs causes defects in both actin assembly at and organization of LC3- and GABARAP-associated structures.

**Figure 5.**
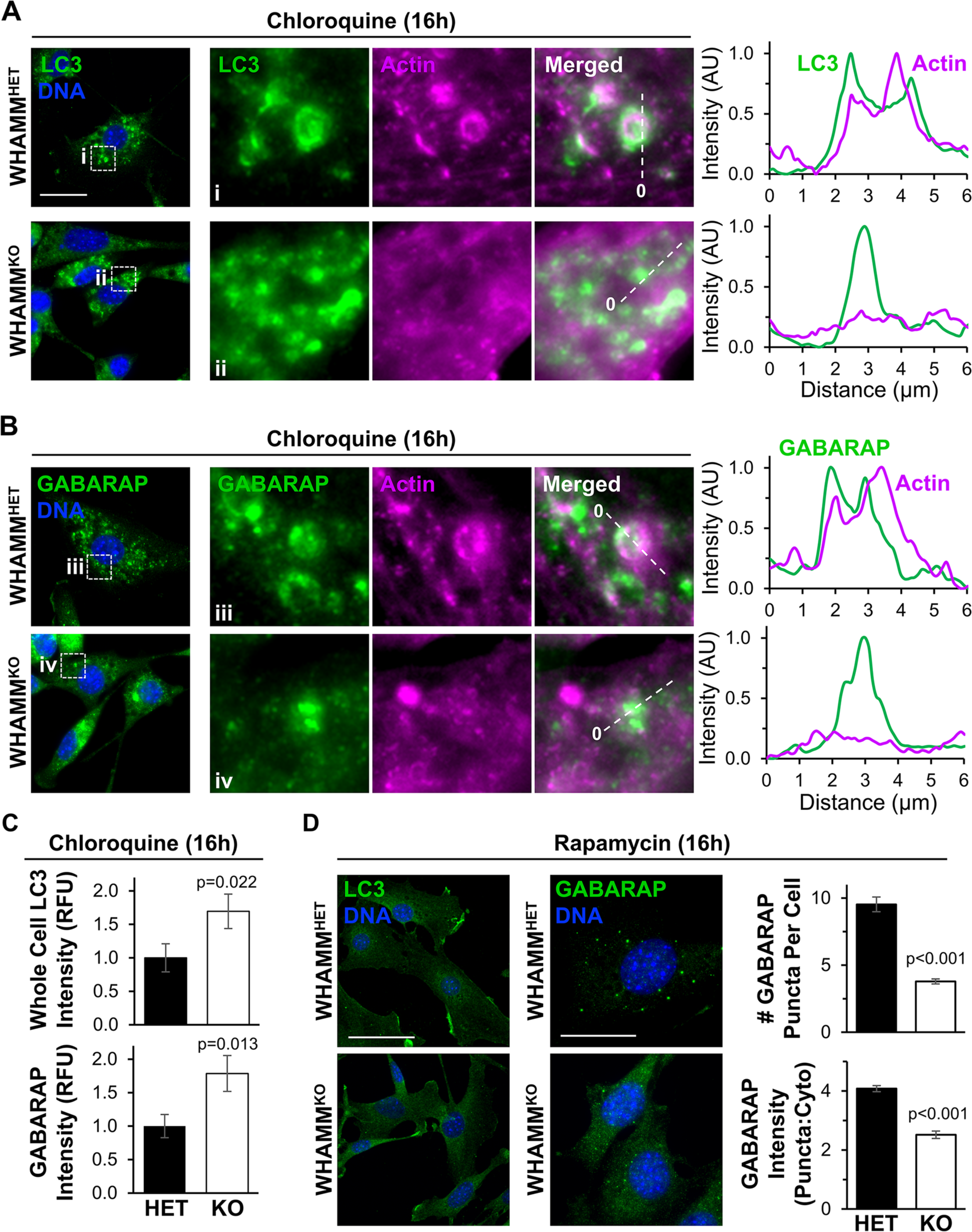
Autophagosome organization and actin recruitment are altered in WHAMM-deficient fibroblasts. **(A-B)** Male heterozygous (WHAMM^HET^) and WHAMM knockout (WHAMM^KO^) mouse embryonic fibroblasts (MEFs) were treated with chloroquine for 16h before being fixed and stained with LC3 antibodies (green), an actin antibody (magenta), and DAPI (DNA; blue). Scale bar, 25µm. Magnifications highlight areas of actin recruitment to LC3- or GABARAP-positive structures in WHAMM^HET^ cells (i) and a lack of actin enrichment at autophagosomal puncta in WHAMM^KO^ cells (ii). Lines were drawn through the images to measure pixel intensity profiles. The origin of each line is indicated with a 0. **(C)** Mean LC3 and GABARAP fluorescence values per cell were measured in ImageJ. Each bar represents the mean ±SD from n=3 experiments (145-161 cells per bar). **(D)** WHAMM^HET^ and WHAMM^KO^ MEFs were treated with rapamycin for 16h before being fixed and stained with LC3 or GABARAP antibodies (green) and DAPI (DNA; blue). Scale bars, 50µm, 25µm. The # of GABARAP puncta per cell was counted manually. Each bar represents the mean ±SD from n=3 experiments (30 cells per genotype per experiment). To calculate the puncta:cytoplasmic ratio of GABARAP fluorescence intensities, the mean fluorescence of GABARAP puncta was divided by the mean cytoplasmic GABARAP fluorescence in ImageJ. Each bar represents the mean ±SD from n=3 experiments (4-7 puncta per cell; 6-8 cells per genotype per experiment). Significant p-values are noted (unpaired t-tests).

To determine if increasing the initiation of autophagy could also influence autophagic membrane morphology differentially in WHAMM-proficient versus WHAMM-deficient cells, we exposed MEFs to the autophagy-inducing mTOR inhibitor rapamycin. While neither WHAMM^HET^ nor WHAMM^KO^ cells displayed any discernible LC3-positive autophagosomes after rapamycin treatment, WHAMM^HET^ cells formed several GABARAP puncta (Figure 5D). Quantification of the number of GABARAP puncta per cell and the puncta-to-cytoplasmic GABARAP intensity ratio revealed that, compared to WHAMM^HET^ MEFs, WHAMM^KO^ MEFs did not effectively shape GABARAP-associated autophagic membranes into discrete puncta (Figure 5D). Overall, the presence of disorganized LC3 and/or GABARAP structures in WHAMM^KO^ MEFs demonstrates the importance of WHAMM in promoting autophagic membrane morphogenesis.

### WHAMM activates the Arp2/3 complex to promote actin assembly, proper LC3 organization, and cargo sequestration in proximal tubule cells

While the experiments in MEFs add to the existing literature about WHAMM in autophagy, our *in vivo* studies point to the kidney proximal tubule as the tissue type in which WHAMM function is most crucial. Therefore, we next studied autophagosome morphogenesis in the HK-2 human proximal tubule cell line. To examine the effects of transient WHAMM depletion, we transfected HK-2 cells with control siRNAs or two independent siRNAs targeting the WHAMM transcript.

Immunoblotting of cell extracts verified WHAMM protein knockdowns (Figure 6A). Following treatment of transfected HK-2 cells with chloroquine, control cells displayed LC3-positive circular autophagic structures that were often associated with actin (Figure 6B). In contrast to these cells, but akin to the changes in LC3 morphology and actin assembly observed in WHAMM^KO^ MEFs, WHAMM-depleted HK-2 cells exhibited disorganized LC3 staining and diffuse actin staining (Figure 6B). Linescans of LC3 structures in control cells showed strong actin recruitment at (Figure 6C, i), around (Figure 6C, ii), and within (Figure 6C, iii; Supplemental Figure S8) autophagic membrane vesicles, whereas WHAMM-depleted cells showed disorganized LC3 staining with little actin enrichment (Figure 6C). Thus, permanent WHAMM deletion in fibroblasts and transient WHAMM depletion in proximal tubule cells both hinder autophagic membrane morphogenesis.

**Figure 6.**
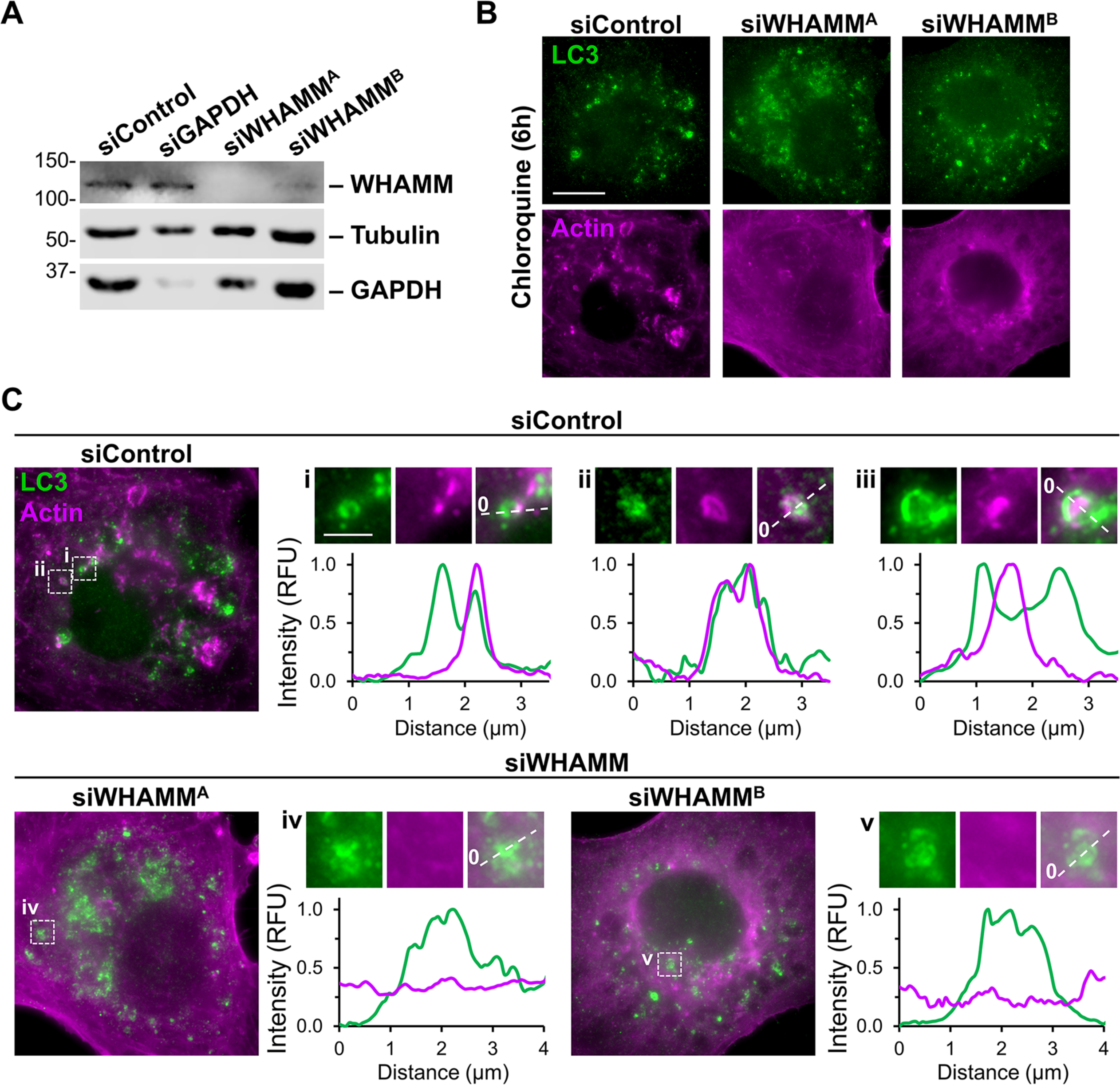
WHAMM depletion disrupts LC3 and actin organization in proximal tubule cells. **(A)** Human kidney proximal tubule (HK-2) cells were transfected with control siRNAs, GAPDH siRNAs, or independent siRNAs targeting the WHAMM transcript before immunoblotting with antibodies to WHAMM, tubulin, and GAPDH. **(B)** Transfected HK-2 cells were exposed to media containing chloroquine for 6h before being fixed and stained with antibodies to LC3 (green) and actin (magenta). Scale bar, 10µm. **(C)** Lines were drawn through the magnified images from (B) to measure pixel intensity profiles. Scale bar, 2µm.

Because the best-characterized molecular activity of WHAMM is to promote Arp2/3 complex-mediated actin polymerization, we next asked whether the Arp2/3 complex impacted autophagosome abundance, localization, or morphology in HK-2 cells. We exposed cells to normal media or to media containing the pharmacological Arp2/3 inhibitor CK666, chloroquine, or CK666 plus chloroquine. Fluorescence microscopy revealed that at steady state, both control and CK666-treated cells exhibited diffuse LC3 staining with no noticeable mature autophagic structures (Figure 7A). Upon lysosomal inhibition with chloroquine, LC3-positive rings accumulated throughout the cytoplasm and often associated with actin (Figure 7B). The combination of chloroquine and CK666 also caused LC3-associated structures to accumulate, but in a smaller perinuclear region, with less circular morphologies, and with minimal recruitment of actin (Figure 7B). Quantification indicated that Arp2/3 inhibition significantly reduced the proportion of cells with mature, ring-shaped LC3-positive autophagosomes, the number of LC3 rings per cell, and the amount of actin localizing at or near the autophagic membrane (Figure 7, C and D). Hence, like WHAMM, the Arp2/3 complex is important for the proper formation of mature, properly-shaped autophagosomes.

**Figure 7.**
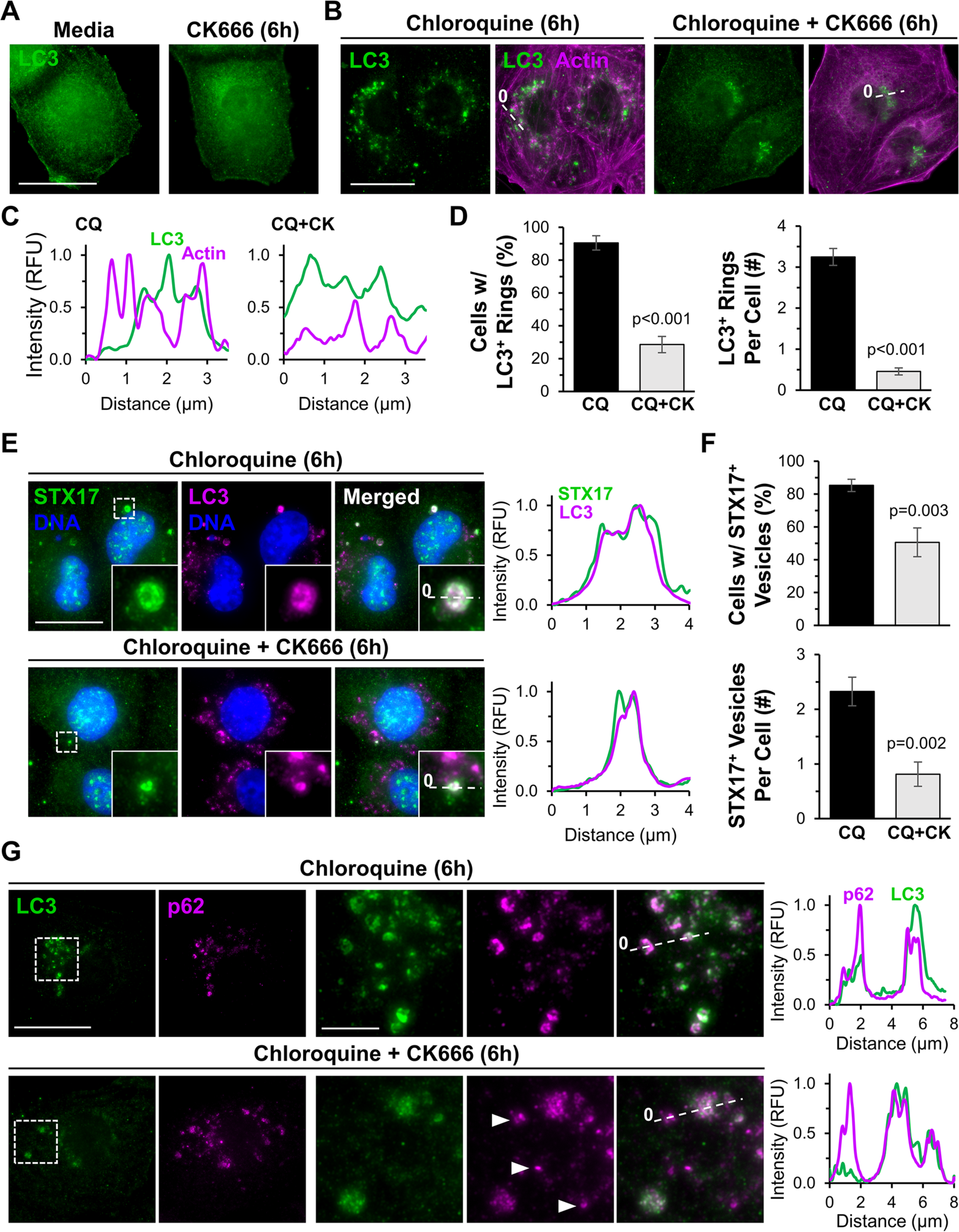
The Arp2/3 complex is crucial for autophagosome closure and actin recruitment during selective autophagy. **(A)** HK-2 cells were treated with normal media or media containing CK666 for 6h before being fixed and stained with LC3 antibodies. **(B)** Cells were treated with chloroquine, or chloroquine plus CK666 for 6h before being fixed and stained with antibodies to LC3 (green) and actin (magenta). **(C)** Lines were drawn through the images in (B) to measure pixel intensity profiles. **(D)** The % of cells with LC3-positive rings and the # of LC3-positive rings per cell were quantified. Each bar represents the mean ±SD from n=3 experiments (150 cells per bar). **(E)** HK-2 cells were treated with chloroquine, or chloroquine plus CK666 for 6h before being fixed and stained with STX17 antibodies (green), an LC3 antibody (magenta), and DAPI (DNA; blue). Scale bar, 20µm. Lines were drawn through magnified images to measure pixel intensity profiles. **(F)** The % of cells with STX17-positive vesicles and the # of STX17-positive vesicles per cell were quantified. Each bar represents the mean ±SD from n=3 experiments (230 cells per bar). **(G)** Cells were treated with media containing chloroquine or chloroquine plus CK666 before being fixed and stained with antibodies to LC3 (green) and p62 (magenta). Arrowheads highlight LC3-independent p62 structures. Lines were drawn through the images to measure pixel intensity profiles. Significant p-values are noted (unpaired t-tests).

If the irregular morphologies of LC3-associated membranes are incompatible with efficient autophagosome closure, then the localization of proteins involved in autophagosome-lysosome fusion should be altered. Syntaxin-17 (STX17) is a SNARE protein that mediates autophagosome closure and is important for subsequent lysosome fusion (Itakura *et al*., 2012; Tsuboyama *et al*., 2016). To examine STX17 localization in the presence and absence of Arp2/3 complex activity, we treated HK-2 cells with DMSO or CK666, inhibited lysosomes with chloroquine, and stained the cells with antibodies to STX17 and LC3 (Figure 7E). Over 80% of DMSO-treated cells possessed STX17-positive vesicles, and on average each cell had two such vesicles (Figure 7E and F). Arp2/3 inhibition reduced both the percentage of cells containing STX17-positive vesicles and the number of STX17-associated vesicles per cell, while the vesicles that were present appeared smaller (Figure 7E and F). These findings further support the conclusion that efficient autophagosome closure is reliant on the Arp2/3 complex.

In the canonical selective autophagy pathway, LC3-decorated membranes interact with autophagy receptors, which are physically linked to ubiquitinated cargo (Vargas *et al*., 2023). So we next asked whether the steps of receptor-ubiquitin or receptor-LC3 interactions were influenced by the Arp2/3 complex. SQSTM1/p62, an autophagy receptor with a wide range of selective targets, has been shown to associate with ubiquitinated protein aggregates that are destined for autophagic degradation (Pankiv *et al*., 2007; Sarraf *et al*., 2020). We therefore induced the production of truncated proteins in HK-2 cells using puromycin and assessed the effects of Arp2/3 inhibition on the recruitment of p62 to ubiquitinated material. Puromycin caused the formation of bright foci of ubiquitinated proteins, and p62 localized to the ubiquitin foci in the absence or presence of CK666 (Supplemental Figure S9), indicating that Arp2/3 inactivation does not prevent the association between ubiquitinated cargo and p62. To evaluate the effect of Arp2/3 inhibition on the step of receptor-LC3 engagement, we examined the recruitment of p62 to LC3. As expected, p62 localized to LC3-positive ring-shaped autophagosomes when cells were treated with chloroquine (Figure 7G). In cells subjected to concurrent chloroquine and CK666 treatment, the disorganized LC3 structures often co-stained for p62 (Figure 7G). However, the Arp2/3-inhibited cells contained additional p62 puncta that were independent of LC3 (Figure 7G). These results suggest that autophagosomal membrane remodeling is coupled to cargo capture and that this process is inefficient without an active Arp2/3 complex.

To connect the observations that both WHAMM and the Arp2/3 complex impact actin assembly at autophagic membranes, we studied actin organization in HK-2 cells expressing a LAP (localization and affinity purification) tagged version of either wild type WHAMM or a mutant WHAMM lacking a critical Arp2/3-binding residue (WHAMM W807A) (Campellone *et al*., 2008). As predicted, wild type WHAMM localized to LC3-positive structures, and actin was enriched around these autophagic membranes (Figure 8A). Similar to cells depleted of WHAMM (Figure 6), cells expressing the WHAMM W807A mutant protein displayed more diffuse cytoplasmic LC3 staining (Figure 8A). In cases where small LC3 puncta were observed, the puncta associated with WHAMM W807A but lacked actin (Figure 8A). Together, these results show that WHAMM stimulates Arp2/3 complex-mediated actin assembly to control autophagic membrane morphogenesis and cargo sequestration.

**Figure 8.**
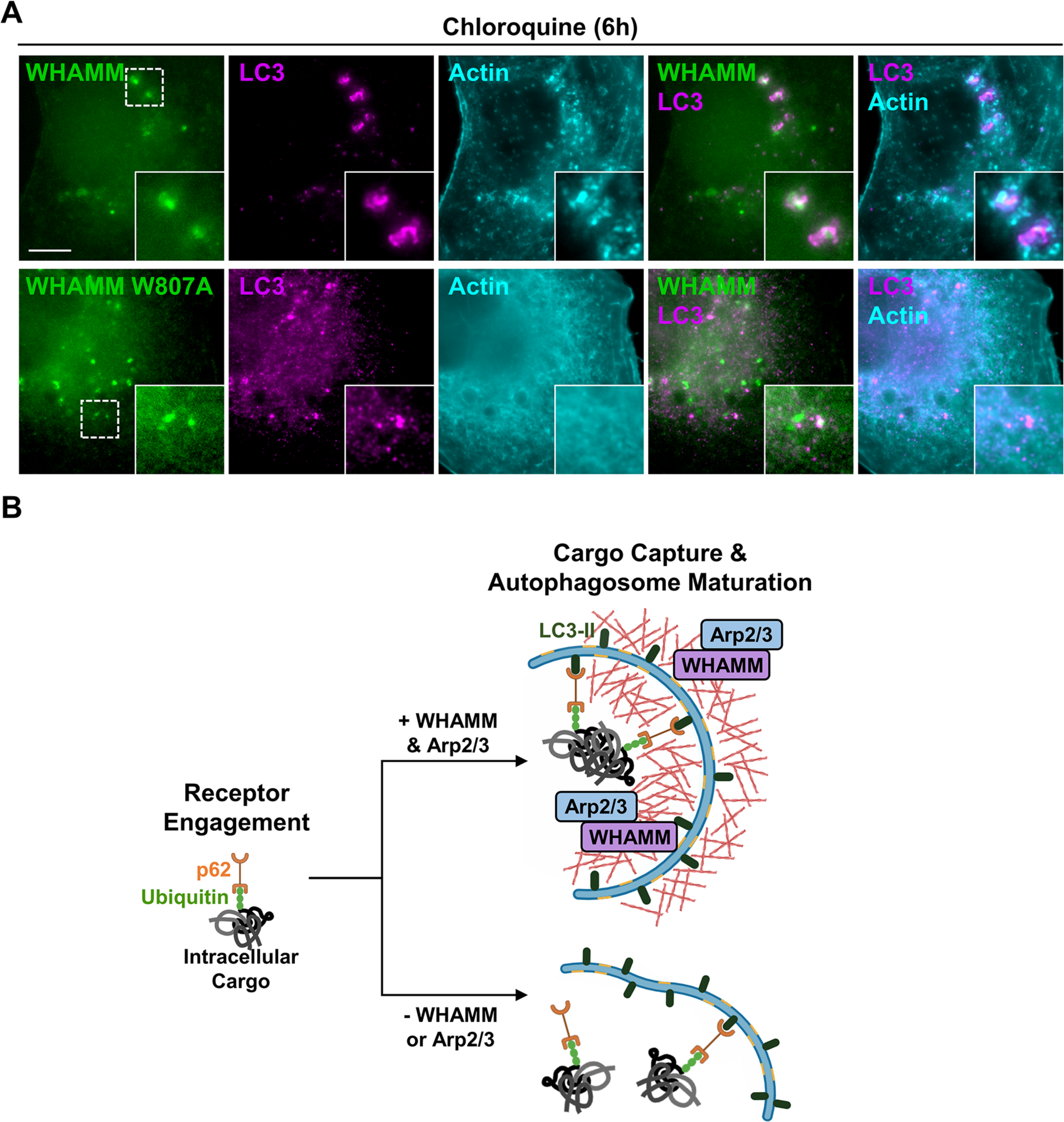
WHAMM promotes Arp2/3 complex-mediated actin assembly during autophagosomal membrane morphogenesis. **(A)** HK-2 cells expressing LAP-WHAMM wild type or a LAP-WHAMM W807A mutant deficient in Arp2/3 activation (both in green) were fixed and stained with antibodies to LC3 (magenta) and actin (cyan). Scale bar, 10µm. **(B)** Model for WHAMM and Arp2/3 complex function in autophagosome closure during cargo capture. Aggregated protein cargo (gray) is ubiquitinated (green) prior to its engagement by autophagy receptors like p62 (orange). WHAMM (purple) and the Arp2/3 complex (blue) promote the assembly of actin filaments (red) necessary for autophagosome membrane morphogenesis. In the absence of WHAMM or the Arp2/3 complex, actin assembly at autophagic membranes is abrogated, resulting in inefficient receptor engagement and autophagosome closure.

## DISCUSSION

Actin nucleation factors are important players in a variety of processes that are crucial for cellular function. However, their roles in organismal health and disease have been relatively understudied. Deletion of several WASP-family members, including N-WASP, WAVE2, and WASH, results in embryonic lethality in mice (Snapper *et al*., 2001; Yan *et al*., 2003; Gomez *et al*., 2012; Xia *et al*., 2013), demonstrating the essentiality of these factors in mammalian development. Similar animal studies of WASP-family proteins from the WHAMM/JMY subgroup had not been previously explored, so we focused our investigation on the impact of WHAMM inactivation in mice. Here we establish the importance of WHAMM in kidney physiology *in vivo* and in autophagosome closure during cargo capture in proximal tubule cells.

The inherited neurodevelopmental and kidney disorder Galloway-Mowat Syndrome is associated with several different *WDR73* mutations (Colin *et al*., 2014; Ben-Omran *et al*., 2015; Vodopiutz *et al*., 2015; Jiang *et al*., 2017; El Younsi *et al*., 2019; Tilley *et al*., 2021), but in patients from Amish communities, 26 of 27 affected individuals were found to be doubly homozygous for loss-of-function mutations in both *WDR73* and *WHAMM*, with one *WHAMM* heterozygous individual presenting with neurological symptoms but lacking renal symptoms (Jinks *et al*., 2015; Mathiowetz *et al*., 2017). Given the genotypic and phenotypic variability of GMS, the extent to which WHAMM inactivation might modify the clinical outcomes in Amish patients has been unclear. Our current study begins to untangle the complexities of Amish GMS by showing that a targeted mutation in *WHAMM* by itself can cause kidney dysfunction in mice.

An elevated urinary protein:creatinine ratio and end-stage renal disease affect the majority of Amish GMS patients (Jinks *et al*., 2015). Recently, a targeted deletion in mouse *Wdr73* was found to be embryonic lethal (Li *et al*., 2022). However, a conditional *Wdr73* deletion in podocytes, terminally differentiated cells required for filtration in the glomerulus, yielded live mice. Such animals did not have any detectable kidney phenotypes until chemically-induced glomerular injury caused albuminuria (Li *et al*., 2022). These results contrast the loss of WHAMM in mice, which is not lethal, but results in male-specific excretion of a low molecular weight protein, multiple solutes, and amino acids in the urine. Thus, while the molecular and cellular basis of GMS pathogenesis still requires much investigation, our characterization of kidney abnormalities in WHAMM-deficient mice supports the idea that the Amish *WHAMM* mutation may be a modifier in the nephrotic aspects of GMS.

Earlier studies in mice indicated that the WASP-family member N-WASP is also important for kidney function (Schell *et al*., 2013; Schell *et al*., 2018). N-WASP regulates Arp2/3- and actin-driven membrane protrusions in podocytes, and deletion of Arp3 increases urinary excretion of albumin (Schell *et al*., 2018). Our current studies with WHAMM knockout mice provide new evidence that the WASP family also contributes to kidney physiology in a different anatomical location, the proximal tubule, which is crucial for reabsorption of small molecules from the filtrate.

The excretion profiles observed in WHAMM-deficient mice are reminiscent of those in other proximal tubule disorders, collectively referred to as Fanconi Syndromes (van der Wijst *et al*., 2019; Lemaire, 2021). Dent disease type II (Hoopes *et al*., 2005; Utsch *et al*., 2006) and Lowe Syndrome (Zhang *et al*., 1995; Bockenhauer *et al*., 2008; Mehta *et al*., 2014) are X-linked Fanconi disorders caused by mutations in the *OCRL* gene. *OCRL* encodes a lipid phosphatase, and loss of its enzymatic activity results in the accumulation of PI(4,5)P2, which leads to defects in endosomal trafficking and autophagy (Vicinanza *et al*., 2011; De Leo *et al*., 2016; Daste *et al*., 2017; De Matteis *et al*., 2017). Given the endocytic and receptor degradation anomalies identified in Lowe Syndrome cells, we tested whether the levels of reabsorption receptors were altered in WHAMM^KO^ kidneys but found no gross changes in receptor abundance. However, the irregular distribution of ACE2 staining in WHAMM knockout tissue is suggestive of defects in proximal tubule organization, polarity, or membrane trafficking. Interestingly, alterations in the urinary proteome, proximal tubule reabsorption, and ACE2 localization have also been observed upon disruption of Vps34, a PI-3 kinase with critical roles in autophagosome biogenesis and endocytic trafficking (Grieco *et al*., 2018; Rinschen *et al*., 2022).

WHAMM was first studied for its functions in microtubule binding, anterograde transport, and Golgi morphogenesis (Campellone *et al*., 2008), but stainings of WHAMM^KO^ MEFs indicated that microtubule and Golgi organization were relatively normal. Taken together with the observations that Amish GMS patient cells have more dramatic defects in autophagosomes than secretory organelles (Mathiowetz *et al*., 2017), we focused our attention on autophagy.

Previous localization and loss-of-function approaches in human epithelial cells, fibroblasts, monkey kidney cells, and GMS patient samples demonstrated the importance of WHAMM early in autophagy during the steps of autophagosome biogenesis, enlargement, and movement (Kast *et al*., 2015; Mathiowetz *et al*., 2017). Additional work in human and rat kidney cell lines showed that WHAMM also plays a role later in autophagy by promoting autolysosome tubulation and turnover (Dai *et al*., 2019; Wu *et al*., 2021).

Our current study evaluated several autophagy-related parameters in mouse kidneys, mouse fibroblasts, and human proximal tubule cells to link the physiological dysfunction *in vivo* with alterations in autophagy *in vitro*. Higher levels of lipidated LC3 in male WHAMM^KO^ kidney tissue first suggested that changes in autophagy accompanied the deficiencies in reabsorption. Fibroblasts generated from WHAMM-proficient or knockout mouse embryos next demonstrated that permanent WHAMM deletion not only resulted in higher overall cellular levels of the LC3 and GABARAP classes of ATG8-family proteins, but also caused aberrations in autophagosomal membrane morphology and a reduction in actin recruitment. In proximal tubule cells, WHAMM and its actin nucleating binding partner, the Arp2/3 complex, were also crucial for the proper morphogenesis and closure of LC3-associated autophagosomes. These results give rise to a model in which WHAMM activates Arp2/3-mediated actin assembly to shape autophagosomal membranes in such a way that allows them to effectively capture autophagy receptors like p62 (Figure 8B). Without WHAMM or the Arp2/3 complex, receptor sequestration becomes inefficient, and the incompletely-sealed misshapen LC3-associated membranes accumulate because their subsequent fusion with lysosomes is impaired.

The numerous functions of WHAMM throughout the autophagy pathway are likely coordinated based on the expression profiles of other proteins that regulate the initiation and progression of multiple degradation pathways. Genetically programmed, epigenetically modulated, or even stochastic changes in such regulatory factors probably determine when and where WHAMM function is most important in a particular cell type. Moreover, different compensatory changes that occur when WHAMM is permanently deleted versus transiently depleted add an additional layer of complexity to deciphering its usual cellular responsibilities. The complicated nature of mammalian protein homeostasis systems and their many potential connections to the actin assembly machinery are underscored by diversity in the repertoires of the (6) LC3/GABARAP isoforms and the (5) basic autophagy receptors (Lazarou *et al*., 2015; Nguyen *et al*., 2016), and perhaps even in the WASP-family members (e.g., WHAMM and JMY) themselves. In the case of mice housed in standard laboratory conditions, the crucial functions for WHAMM appeared anatomically in the kidney proximal tubule and molecularly in LC3/GABARAP-associated autophagosome morphogenesis and closure.

The distinct excretion phenotypes in male versus female WHAMM^KO^ mice, and the fact that tissue from only male knockouts showed substantial changes in LC3 lipidation, highlight a relationship between alterations in kidney function and molecular malfunctions in autophagy. The specific reasons for this male specificity are unclear, but other sex-specific discrepancies *in vivo* have been attributed to differences in the distribution and/or abundance of transporters throughout the nephron (Veiras *et al*., 2017; Harris *et al*., 2018; Li *et al*., 2018; Hu *et al*., 2020; Torres-Pinzon *et al*., 2021). Anatomically, males have a greater density of proximal tubules in the cortex, whereas in females the collecting duct comprises a larger volume (Harris *et al*., 2018). Modeling further suggests that a smaller transport area and varied expression of transporters results in less tubular reabsorption in females (Li *et al*., 2018). Autophagy rates also differ between males and females (Shang *et al*., 2021). Future work on the mechanisms regulating autophagy in the proximal tubule is required to better understand the sex-based differences in kidney physiology arising from the loss of WHAMM.

Lowe Syndrome is X-linked, and excessive actin polymerization on multiple organelles is a prominent feature in cells from patients with this disorder (Suchy and Nussbaum, 2002; Vicinanza *et al*., 2011; Festa *et al*., 2019; Berquez *et al*., 2020). Alterations in PI(3)P and PI(4,5)P2 membrane composition are key drivers of signaling to the actin nucleation machinery in this context, and N-WASP is at least partly responsible for the ectopic actin assembly (Vicinanza *et al*., 2011; Daste *et al*., 2017). However, WHAMM is also capable of binding to PI(3)P (Mathiowetz *et al*., 2017) and PI(4,5)P2 (Dai *et al*., 2019). So on one hand it is tempting to speculate that WHAMM-mediated Arp2/3 activation may influence the excess cytoskeletal rearrangements and trafficking modifications that take place in Lowe Syndrome cells, while on the other hand, reduced actin assembly may underlie the autophagy defects in WHAMM- depleted cells. This combination of previous data and our new findings are consistent with the idea that either over-or under-active actin assembly pathways can cause similar tubular reabsorption problems. Thus, proximal tubule function is governed by exquisitely tight spatiotemporal control of phospholipid signaling to the actin polymerization machinery. Future characterizations of the connections between key autophagy regulators and actin nucleation factors therefore hold promise for determining how proteostasis pathways and cytoskeletal activities collaborate in healthy kidneys and how their functions are altered during distinct diseases.

## MATERIALS AND METHODS

### Mice

A floxed *Whamm* allele generated by the Knockout Mouse Project (genome.gov/17515708) incorporated SA-IRES-*lacZ*-pA and *neo* cassettes between *Whamm* exons 2 and 3, with *loxP* sites located between the two cassettes and following exon 3. Cre-mediated recombination using the B6.C-Tg(CMV-Cre)1Cgn/J strain resulted in the excision of *neo* and exon 3, giving rise to a *Whamm* knockout allele. These B6N(Cg)-*Whamm*^tm1b(KOMP)Wtsi^/3J mice were obtained from The Jackson Laboratory (JAX stock JR#027472). Animals were maintained on pine shavings and given a standard rodent diet (LabDiet 5KOG) and acidified water in a pathogen-free room that was maintained at 21°C with a 12h light/dark cycle (6am to 6pm). Experimental animals were generated by mating heterozygous knockout (WHAMM^HET^) mice and selecting homozygous knockout (WHAMM^KO^) offspring and their wild type (WHAMM^WT^) littermates. Genotyping primers were designed to amplify a 221bp fragment of the wild type *Whamm* allele or a 222bp fragment of the knockout *Whamm* allele (Supplemental Table S1). All mice were sacrificed during the SARS-CoV-2 pandemic in 2020. Sperm is cryopreserved at JAX.

### Mouse Phenotyping

Spot urine and blood were collected at 8, 16, 20, and 24 weeks-of-age. Urinary albumin, glucose, phosphate, potassium, sodium, chloride, calcium, and creatinine, and plasma glucose were measured using a Beckman Coulter DxC 700 AU chemistry analyzer. For relative quantification of amino acids in the urine, a LC-MS/MS Selected Reaction Monitoring (SRM) method was performed using a Thermo Fisher Scientific TSQ Endura equipped with a Vanquish UPLC based on Thermo Fisher’s technical note 65382 adapted for smaller volumes of urine.

Briefly, SRM transitions were detailed in the TraceFinder software and then verified with the Metabolomics Amino Acid Mix Standard (Cambridge Isotope Laboratories) for unique transitions. 10μL of urine was precipitated with 30% sulfosalicylic acid (final concentration 10%) and vortexed for 30s. Samples were allowed to precipitate for 30min at 4°C and centrifuged to pellet protein. Supernatant was then mixed with internal standard and diluent mixture. 4μL of this final solution was injected into the platform. Chromatographic separation was performed over 18min, with an Acclaim Trinity mixed mode column. Buffer A was ammonium formate in water, pH ∼3. Buffer B was acetonitrile with ammonium formate, pH ∼3. Separation was achieved with a two-part separation and flow rate increase. Detection of each of the 52 transitions was performed with the TSQ Endura triple quadruple mass spectrometer. Data was acquired in SRM mode using a resolution of 0.7 m/z full width at half maximum with a 500ms cycle time. Data were processed using Tracefinder 4.1 software.

### Bioinformatics

Gene expression maps from adult mice were created using Kidney Cell Explorer (Ransick *et al*., 2019). This searchable database (<u>cello.shinyapps.io/kidneycellexplorer/</u>), consisting of single cell RNA-sequencing data clustered into distinct anatomical regions (“metacells”) of the nephron, was used to generate heat maps of the normalized average expression of genes (average expression) as well as the proportion of cells expressing a gene (expressed proportion) within each metacell.

### Kidney Histology and Immunostaining

For histological analyses, kidneys were collected from 25-week-old mice in 10% neutral buffered formalin, embedded in paraffin, and subjected to periodic acid-Schiff (PAS) staining. Histological slides can be viewed at https://images.jax.org/webclient/?show=dataset-2763. For immunostaining, 5µm kidney sections were mounted onto charged glass slides, deparaffinized in Histo-Clear (National Diagnostics) twice, 100% ethanol twice, 95% ethanol twice, and 70% ethanol, then rehydrated in deionized water. Antigen retrieval was performed at 95°C in citrate buffer (20mM citric acid, 93mM sodium citrate in water) pH 6.0 for 30min, and slides were cooled to room temperature before 3 phosphate-buffered saline (PBS) washes. Tissue samples were incubated in blocking buffer (PBS containing 1% bovine serum albumin (BSA), 10% goat serum, 0.1% Tween-20) for 2h at room temperature, washed once with PBS, probed with primary antibodies (Supplemental Table S2) for 10h at 4°C, washed 3 times, and treated with AlexaFluor-conjugated secondary antibodies and DAPI (Supplemental Table S2) for 2h at 4°C, followed by 3 PBS washes and mounting using ProLong Gold (Invitrogen) and 18mm square glass coverslips. Slides were imaged as described below.

### Kidney Tissue Preparation

Kidneys were harvested from 25-week-old WHAMM^WT^ and WHAMM^KO^ mice after cervical dislocation and frozen at ™80°C. Tissue extracts were prepared by resuspending thawed kidney thirds in tissue lysis buffer (20mM HEPES pH 7.4, 100mM NaCl, 1% IGEPAL CA-630, 1mM EDTA, 1mM Na3VO4, 1mM NaF, plus 1mM PMSF, and 10μg/ml each of aprotinin, leupeptin, pepstatin, and chymostatin) and sonicating at 60% power for 35s 3 times using a Fisher dismembranator. The lysates were then clarified by centrifugation at 21,000xg for 12min at 4°C, and the supernatants were collected and centrifuged again at 21,000xg for 6min at 4°C. Extract concentrations were measured using Bradford assays (Bio-Rad), aliquoted, and stored at ™80°C. **Cell Culture** To isolate MEFs, timed matings between WHAMM^HET^ mice were performed and embryos collected at E13.5 in Dulbecco’s Modified Eagle Medium (DMEM) containing L-glutamine. Embryos were shipped on ice overnight and within 24h of harvesting, the livers, hearts, and brains were removed, and the remaining tissues were manually dissociated with scalpels. Embryonic slurries were each transferred into 3mL of 0.25% Trypsin-EDTA and incubated on ice for 18h. Without disturbing the settled tissue, 2mL of the supernatants were removed, and the remaining materials were incubated at 37°C for 30min. Each cell suspension was mixed with 9mL of complete media (DMEM containing 10% fetal bovine serum (FBS), GlutaMax, and antibiotic-antimycotic (Gibco)) and allowed to adhere to a 10cm dish at 37°C in 5% CO2 for 24h. Adherent cells were washed with PBS, collected in 0.05% Trypsin-EDTA, resuspended in media, and split into two new 10cm dishes. After an additional 24h of growth, the cells in one dish were washed with PBS, collected in PBS containing 2mM EDTA, pelleted, resuspended in FBS containing 10% DMSO, aliquoted, and stored in liquid nitrogen. The cells in the second dish were maintained at 35-95% confluence, passaged every 2-3 days, and cryopreserved after passage 7. Upon reanimation, cells went through crisis after 3-5 more passages, and immortalized cultures eventually emerged. Experiments were conducted after passage 16. Human male HK-2 proximal tubule kidney cells (ATCC) were cultured in DMEM, 10% FBS, GlutaMax, and antibiotic-antimycotic. All experiments were performed using cells that had been in active culture for 2-10 trypsinized passages after thawing.

### Cell Genotyping

MEFs grown in 6cm dishes were collected in PBS containing 1mM EDTA, centrifuged, washed with PBS, and recentrifuged. Genomic DNA was isolated from ∼2×10^6^ cells using the Monarch DNA purification Kit (New England Biolabs). For genotyping, PCRs were performed using 50ng of genomic DNA, gene-specific primers, and Taq polymerase (New England Biolabs). Primers (Supplemental Table S1) were designed to amplify the wild type and knockout alleles as described above, 480bp and 660bp fragments of the *Xlr* gene on the X chromosome, a 280bp fragment of the *Sly* gene on the Y chromosome, and a 241bp *Gapdh* control. PCR products were subjected to ethidium bromide agarose gel electrophoresis and visualized using ImageJ (Schindelin *et al*., 2012). Male heterozygous (WHAMM^HET^X/Y) and male homozygous *Whamm* knockout (WHAMM^KO^X/Y) cell populations were generated, but we were unable to isolate male wild type MEFs despite multiple attempts.

### Immunoblotting

Cells grown in 6-well plates were collected in PBS containing 1mM EDTA and centrifuged before storing at ™20°C. Pellets were resuspended in cell lysis buffer (20mM HEPES pH 7.4, 50mM NaCl, 0.5mM EDTA, 1% Triton X-100, 1mM Na3VO4, and 1mM NaF, plus protease inhibitors). Kidney or cell extracts were diluted in SDS-PAGE sample buffer, boiled, centrifuged, and subjected to SDS-PAGE before transfer to nitrocellulose (GE Healthcare). Membranes were blocked in PBS containing 5% milk (PBS-M) before being probed with primary antibodies (Supplemental Table S2) diluted in PBS-M overnight at 4°C plus an additional 2-3h at room temperature. Membranes were rinsed twice with PBS and washed thrice with PBS + 0.5% Tween-20 (PBS-T). Membranes were then probed with secondary antibodies conjugated to IRDye-800, IRDye-680, or horseradish peroxidase (Supplemental Table S2), rinsed with PBS, and washed with PBS-T. Blots were visualized using a LI-COR Odyssey Fc imaging system, band intensities determined using the Analysis tool in Image Studio software, and quantities of proteins-of-interest normalized to tubulin, actin, and/or GAPDH loading controls.

### Chemical Treatments and Transfections

Cells were seeded onto 12mm glass coverslips in 24-well plates, allowed to grow for 24h, and then treated prior to fixation. MEFs were treated with media containing 50µM chloroquine (Sigma) or 10µM rapamycin (Tocris) for 16h prior to fixation. HK-2 cells were treated with media containing 50µM chloroquine, 200µM CK666 (Calbiochem), or both for 6h prior to fixation, or with media containing 5µg/mL puromycin (Sigma), or puromycin plus 200µM CK666 for 2h prior to fixation. For RNAi experiments, cells were grown in 6-well plates for 24h, transfected with 40nM siRNAs (Supplemental Table S1) using RNAiMAX (Invitrogen), incubated in growth media for 24h, reseeded onto 12mm glass coverslips, and incubated for an additional 48h prior to fixation. For transgene expression, HK-2 cells grown on 12mm glass coverslips were transfected with 50-100ng of LAP-WHAMM(WT) or LAP-WHAMM(W807A) (Supplemental Table S1) using LipofectamineLTX with Plus reagent (Invitrogen) diluted in DMEM. After 5h, cells were incubated in growth media for an additional 18h prior to fixation. The LAP tag consists of an N-terminal His-EGFP-TEV-S peptide (Campellone *et al*., 2008).

### Immunofluorescence Microscopy

Cells grown on coverslips were fixed using 2.5% paraformaldehyde (PFA) in PBS for 30min, washed, permeabilized with 0.1% TritonX-100 in PBS, washed, and incubated in blocking buffer (PBS containing 1% FBS, 1% BSA, and 0.02% NaN3) for 15min. For LC3 and GABARAP staining, a methanol denaturation step was included between the PFA and TritonX-100 steps. Cells were probed with primary antibodies (Supplemental Table S2) for 45min, washed, and treated with AlexaFluor-conjugated secondary antibodies, DAPI, and/or AlexaFluor-conjugated phalloidin (Supplemental Table S2) for 45min, followed by washes and mounting in ProLong Gold. All tissue and cell images were captured using a Nikon Eclipse T*i* inverted microscope equipped with Plan Apo 100X/1.45, Plan Apo 60X/1.40, or Plan Fluor 20x/0.5 numerical aperture objectives, an Andor Clara-E camera, and a computer running NIS Elements software. Cells were viewed in multiple focal planes, and Z-series were captured in 0.2µm steps. Images presented in the figures represent either 4-5 slice projections for tissue samples or 1-3 slice projections for cell samples.

### Image Processing and Quantification

Images were processed and analyzed using ImageJ (Schindelin *et al*., 2012). For analyses of ACE2 and Megalin fluorescence intensities per tubule, the Selection tool was used to outline proximal tubule cell clusters, the Measure tool was used to acquire their mean fluorescence intensities, and the background signals from outside of the tubules were subtracted from the mean fluorescence values. To calculate the ACE2 polarity ratios, the selection tool was used to select ∼12µm^2^ areas in the apical and cytoplasmic regions of the cluster of proximal tubule cells, the Measure tool was used to acquire the mean fluorescence intensity in each region, background fluorescence was subtracted from these values, and the ACE2 polarity ratio was determined by dividing the apical by cytoplasmic fluorescence per tubule. To generate ACE2 pixel intensity plots, an 8µm line was drawn through a cluster of proximal tubule cells such that the apical region was at the midpoint, and the Plot Profile tool was used to measure the intensity along the line. The background signal from outside of the tubule was subtracted from the mean fluorescence values, and the maximum ACE2 intensity for each wild type profile was set to 1.

For analyses of LC3/GABARAP morphology and actin localization in MEFs and HK-2s, a 3-6µm line was drawn through a fluorescent autophagic structure, such that the center of the circle or punctum was at the midpoint, and the Plot Profile tool was used to measure the intensity along the line. To determine the fluorescence intensities of LC3 and GABARAP, the Selection tool was used to outline individual cells, and the Measure tool was used to acquire total fluorescence. The number of GABARAP puncta per cell was counted manually. To calculate the puncta-to-cytoplasmic ratio of GABARAP fluorescence, the Selection tool was used to select GABARAP puncta or an equivalently-sized area of the cytoplasm, the Measure tool was used to acquire the mean fluorescence intensity in each area, and the puncta-to-cytoplasmic GABARAP fluorescence ratio was determined. The percentage of cells with LC3- positive rings and the number of LC3-positive rings per cell were counted manually. The percentage of cells with STX17-positive vesicles and the number of STX17-positive vesicles per cell were counted manually. Vesicles were confirmed to be autophagic based on LC3 staining.

### Reproducibility and Statistics

For urinalyses, quantifications were based on data from 4-10 mice of a given genotype. For kidney tissue immunoblotting and immunostaining, quantifications were based on data from kidneys from 3-5 mice per genotype. For cell-based assays, conclusions were based on observations made from at least 3 separate experiments, and quantifications were based on data from 3 representative experiments. The sample size used for statistical tests was the number of mice or the number of times an experiment was performed. Statistical analyses were performed using GraphPad Prism software. Statistics for data sets comparing 2 conditions were determined using unpaired t-tests as noted in the Legends. Statistics for data sets with 3 or more conditions were performed using ANOVAs followed by Tukey’s post-hoc test unless otherwise indicated. P-values <0.05 were considered statistically significant.

## ACKNOWLEDGEMENTS

We gratefully acknowledge the contribution of Dorothy Ahlf Wheatcraft and the Protein Sciences and Histopathology Sciences services at The Jackson Laboratory for expert assistance with the work described in this publication. We also thank Naydu Nunno for help processing kidney samples and Campellone Lab members for their comments on this manuscript. KGC was supported by National Institutes of Health grants GM107441 and AG050774 (www.nih.gov). RK was supported by National Institutes of Health grants ES29916, AG038070, DK131019, and DK131061, and the Alport Syndrome Foundation. The funders had no role in study design, data collection and analysis, decision to publish, or preparation of the manuscript.

