## Supplemental Information for "WHAMM functions in kidney reabsorption and polymerizes actin to promote autophagosomal membrane closure and cargo sequestration"

### Table S1. Animals, Cell Lines, Primers, siRNAs, and Plasmids

***Mice***

| Description | Strain | Source |
|---|---|---|
| $WHAMM^{HET}$ | B6N(Cg)-*Whamm*$^{tm1b(KOMP)Wtsi}$/3J | The Jackson Laboratory (JR#027472) |
| $WHAMM^{WT}$ | $WHAMM^{WT}$ | This Study |
| $WHAMM^{KO}$ | $WHAMM^{KO}$ | This Study |

***Cell Lines***

| Cell Type | Genotype | Source |
|---|---|---|
| HK-2 | | ATCC (CRL-2190) |
| MEF HET Male | *Whamm*$^{HET}{}_{X/Y}$ | This Study |
| MEF KO Male | *Whamm*$^{KO}{}_{X/Y}$ | This Study |

***Primers***

| Description | Species | Sequences | |
|---|---|---|---|
| $WHAMM^{WT}$ | Mouse | F | GGGCAAGATACAACACTGAA |
| $WHAMM^{KO}$ | Mouse | F | CTGGATCCGGAATAACTTCG |
| $WHAMM^{WT/KO}$ | Mouse | R | GGCCAGCTTCCTTCTCATAA |
| SX | Mouse | F<br>R | GATGATTTGAGTGGAAATGTGAGGTA<br>CTTATGTTTATAGGCATGCACCATGTA |
| GAPDH | Mouse | F<br>R | GAGGGCTGCAGTCCGTATTT<br>CCTCCCCCTATCAGTTCGGA |

***siRNAs***

| Target | Identifier |
|---|---|
| Control | Ambion AM4611 |
| GAPDH | Ambion AM4624 |
| WHAMM A<br>WHAMM B | Sigma WD07317596<br>Invitrogen HSS151264 |

***Plasmids***

| Description | Vector | Species | Sources |
|---|---|---|---|
| pKC-LAP-C1 | N/A | N/A | Campellone *et al*., 2008 |
| pKC-LAP-WHAMM | pKC-LAP-C1 | Human | Campellone *et al*., 2008 |
| pKC-LAP-WHAMM W807A | pKC-LAP-C1 | Human | Campellone *et al*., 2008 |

| **<u>Table S2.</u> Immunofluorescence and Immunoblotting Reagents** | | | | |
|---|---|---|---|---|
| Target | Probe | | Conc. | Identifier |
| ***Primary Antibodies (Immunofluorescence)*** | | | | |
| ACE2 | anti-ACE2 | Rabbit | 1:1,000 | Proteintech (21115-I-AP) |
| Actin | anti-β-Actin | Mouse | 1:10,000 | Proteintech (66009-I-Ig) |
| AIF | anti-AIF | Rabbit | 1:1,000 | Cell Signaling Technology (5318) |
| GABARAP | anti-GABARAP | Rabbit | 1:750 | Proteintech (18723-I-AP) |
| GM130 | anti-GM130 | Mouse | 1:1,000 | BD Transduction Laboratories (610822) |
| LC3 | anti-LC3 | Rabbit | 1:750 | Proteintech (14600-I-AP) |
| LC3B | anti-LC3B | Mouse | 1:750 | Cell Signaling (83506S) |
| Megalin | anti-LRP2 | Rabbit | 1:1,000 | Proteintech (19700-I-AP) |
| Microtubules | anti-β-Tubulin | Mouse | 1:1,000 | DSHB (E7) |
| p62 | anti-p62 | Mouse | 1:1,000 | BD Transduction Laboratories (610832) |
| p62 | anti-p62/SQSTM1 | Rabbit | 1:1,000 | Proteintech (18420-I-AP) |
| STX17 | anti-Syntaxin 17 | Rabbit | 1:1,000 | Proteintech (17815-I-AP) |
| Ubiquitin | anti-ubiquitin | Mouse | 1:1,000 | EDM Millipore (ST1200) |
| ***Primary Antibodies (Immunoblotting)*** | | | | |
| ACE2 | anti-ACE2 | Rabbit | 1:1,000 | Proteintech (21115-I-AP) |
| Actin | anti-Beta-Actin | Mouse | 1:10,000 | Proteintech (66009-I-Ig) |
| GABARAP | anti-GABARAP | Rabbit | 1:750 | Proteintech (18723-I-AP) |
| GAPDH | anti-GAPDH | Mouse | 1:10,000 | Proteintech (60004-I-Ig) |
| LC3 | anti-LC3 | Rabbit | 1:750 | Proteintech (14600-I-AP) |
| Megalin | anti-LRP2 | Rabbit | 1:1,000 | Proteintech (19700-I-AP) |
| SGLT2 | anti-SGTL2 | Rabbit | 1:1,000 | Proteintech (24654-I-AP) |
| SLC20A1 | anti-SLC20A1 | Rabbit | 1:1,000 | Proteintech (12423-I-AP) |
| Tubulin | anti-β-Tubulin | Mouse | 1:10,000 | DSHB (E7) |
| WHAMM | anti-WHAMM | Rabbit | 1:1,000 | Shen *et al.*, 2012 |
| ***Secondary Antibodies (Immunofluorescence)*** | | | | |
| Mouse IgG | Alexa 488, 555 anti-mouse | Goat | 4 µg/mL | Life Technologies (A11029; A21424) |
| Rabbit IgG | Alexa 488 anti-rabbit | Goat | 4 µg/mL | Life Technologies (A11034) |
| ***Secondary Antibodies (Immunoblotting)*** | | | | |
| Mouse IgG | IRDye 680 anti-Mouse | Donkey | 0.5 µg/mL | LI-COR (926-68022) |
| Rabbit IgG | IRDye 800 anti-Rabbit | Donkey | 0.5 µg/mL | LI-COR (926-32213) |
| Rabbit IgG | HRP anti-Rabbit | Donkey | 1:10,000 | GE Healthcare (NA934V) |
| ***Molecular Probes (Fluorescence)*** | | | | |
| DNA | 4’,6-diamidino-2-phenylindole (DAPI) | | 1 µg/mL | Invitrogen (D1306) |
| F-actin | AlexaFluor 647-Phalloidin | | 0.4 U/mL | Invitrogen (A22287) |

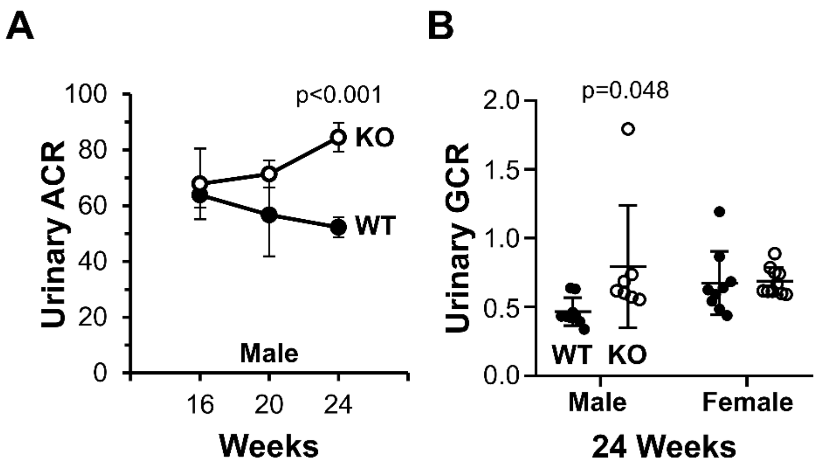

**Figure S1. Male WHAMM$^{KO}$ mice excrete elevated levels of urinary albumin and glucose.** **(A)** Urine samples were collected from male wild type (WT; filled circles) or WHAMM knockout (KO; open circles) mice. Urinary albumin-to-creatinine ratios (ACR) for males from 16 to 24 weeks-of-age were plotted. Each circle represents the mean ±SE from n=6-7 mice per genotype for each timepoint. **(B)** Urinary glucose-to-creatinine ratios (GCR) from male and female mice were plotted. Each circle represents one mouse. Statistical bars display the mean ±SD from n=7-9 mice. Significant p-values are noted (unpaired t-tests).

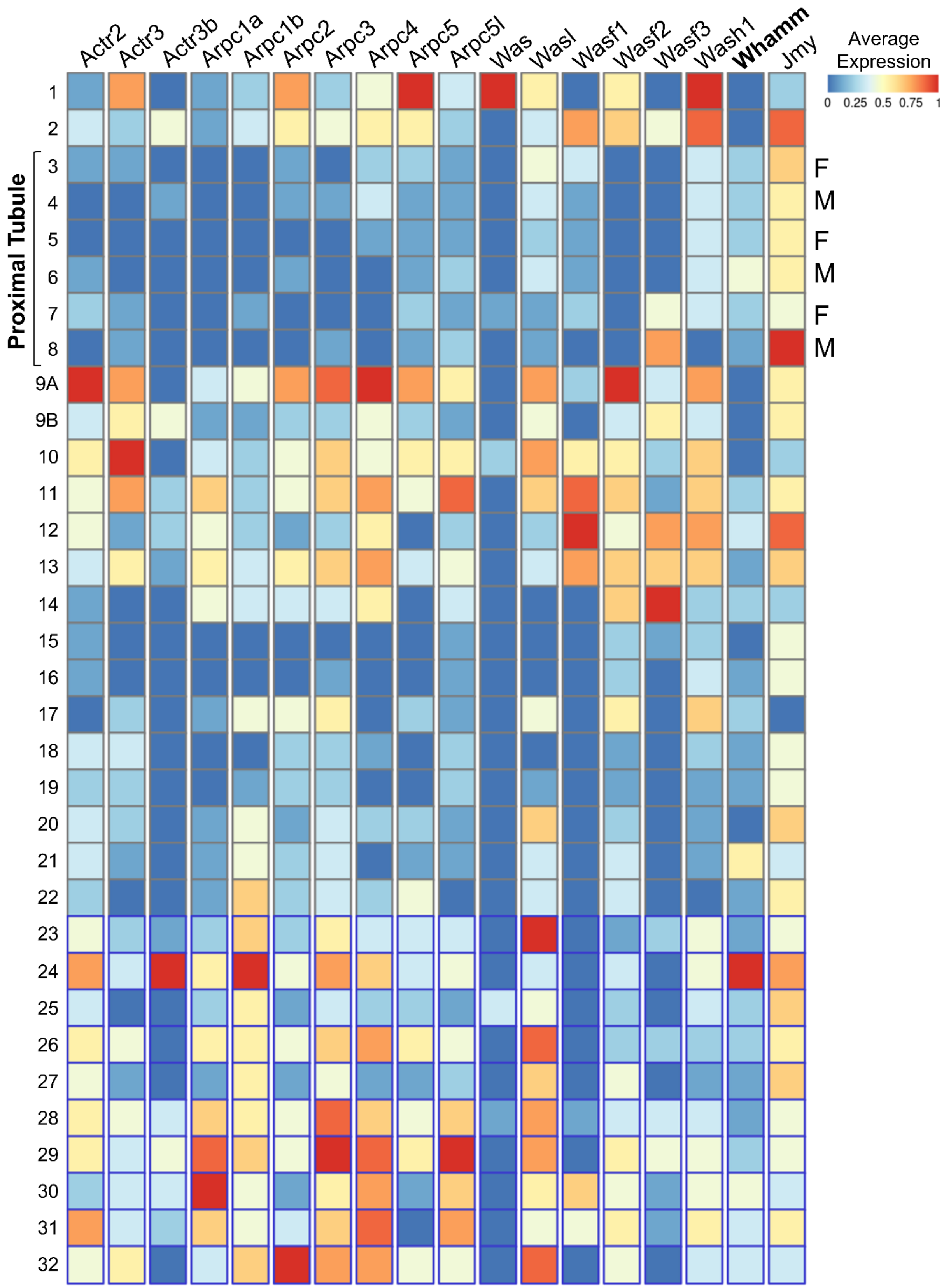
Actr2
Actr3
Actr3b
Arpc1a
Arpc1b
Arpc2
Arpc3
Arpc4
Arpc5
Arpc5l
Was
Wasl
Wasf1
Wasf2
Wasf3
Wash1
Whamm
Jmy
Average Expression
0 0.25 0.5 0.75 1
Proximal Tubule
1
2
3
4
5
6
7
8
9A
9B
F
M
F
M
F
M

**Figure S2. Average relative expression of Arp2/3 complex subunits and WASP-family genes in the nephron.** Data from KidneyCellExplorer (cello.shinyapps.io/kidneycellexplorer; (Ransick *et al.*, 2019)) is based on single-cell RNA sequencing analyses and grouping into "metacell" clusters for mouse nephron regions 1-32. Average expression is plotted on a 0-1 scale and displayed in a heat map, wherein 0 represents no expression for a particular gene and 1 is assigned to the region of the nephron in which that gene's expression is highest. Abbreviations: 1 – Podocytes; 2 – Parietal epithelium; 3 – Segment 1 Proximal Tubule (Female); 4 – Segment 1 Proximal Tubule (Male); 5 – Segment 2 Proximal Tubule (Female); 6 – Segment 2 Proximal Tubule (Male); 7 – Segment 3 Proximal Tubule (Female); 8 – Segment 3 Proximal Tubule (Male); 9A – LOH thin descending limb of inner stripe of outer medulla of cortical nephron; 9B – LOH thin descending limb of inner stripe of outer medulla of juxtamedullary nephron; 10 – Upper LOH thin descending limb of inner medulla of juxtamedullary nephron; 11 – Lower LOH thin descending limb of inner medulla of juxtamedullary nephron; 12 – Lower LOH thin limb of inner medulla of juxtamedullary nephron; 13 – Lower LOH thin limb of inner medulla of juxtamedullary nephron; 14 – Upper LOH thin ascending limb of inner medulla of juxtamedullary nephron; 15 – Distal straight tubule of inner stripe of outer medulla; 16 – Distal straight tubule of outer stripe of outer medulla and cortex; 17 – Macula densa; 18 – Distal convoluted tubule; 19 – Nephron connecting tubule; 20 – Principal-like cell of nephron connecting tubule; 21 – Intercalated type non-A non-B cell of nephron connecting tubule; 22 – Intercalated type A cell of nephron connecting tubule and cortical collecting duct; 23 – Principal-like cell of cortical collecting duct; 24 – Intercalated type B cell of cortical collecting duct; 25 – Intercalated type A cell of outer medullary collecting duct; 26 – Principal cell of outer medullary collecting duct; 27 – Intercalated type A cell of inner medullary collecting duct; 28 – Principal cell of inner medullary collecting duct type 1; 29 – Principal cell of inner medullary collecting duct type 2; 30 – Principal-like cell of deep inner medullary collecting duct type 1; 31 – Cell of deep inner medullary collecting duct type 2; 32 – Deep medullary epithelium of pelvis.

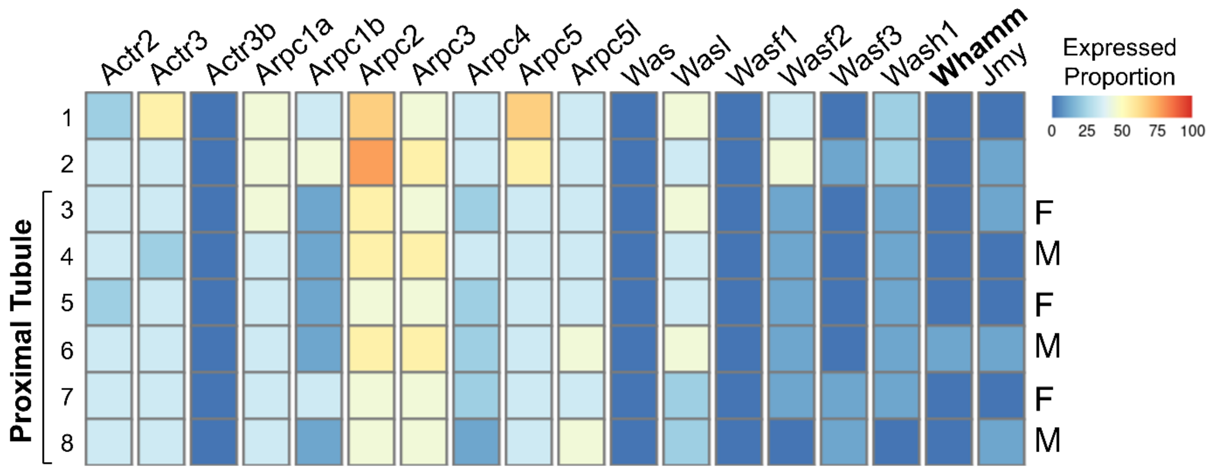

**Figure S3. Proportion of cells expressing Arp2/3 complex subunits and WASP-family genes in the nephron.** Data from KidneyCellExplorer (cello.shinyapps.io/kidneycellexplorer; (Ransick *et al.*, 2019)) is based on single-cell RNA sequencing analyses and grouping into "metacell" clusters for mouse nephron regions 1-32 as in Figure S2. The proportion of cells expressing a gene of interest in each metacell is plotted on a 0-100 scale and displayed according to the heat map.

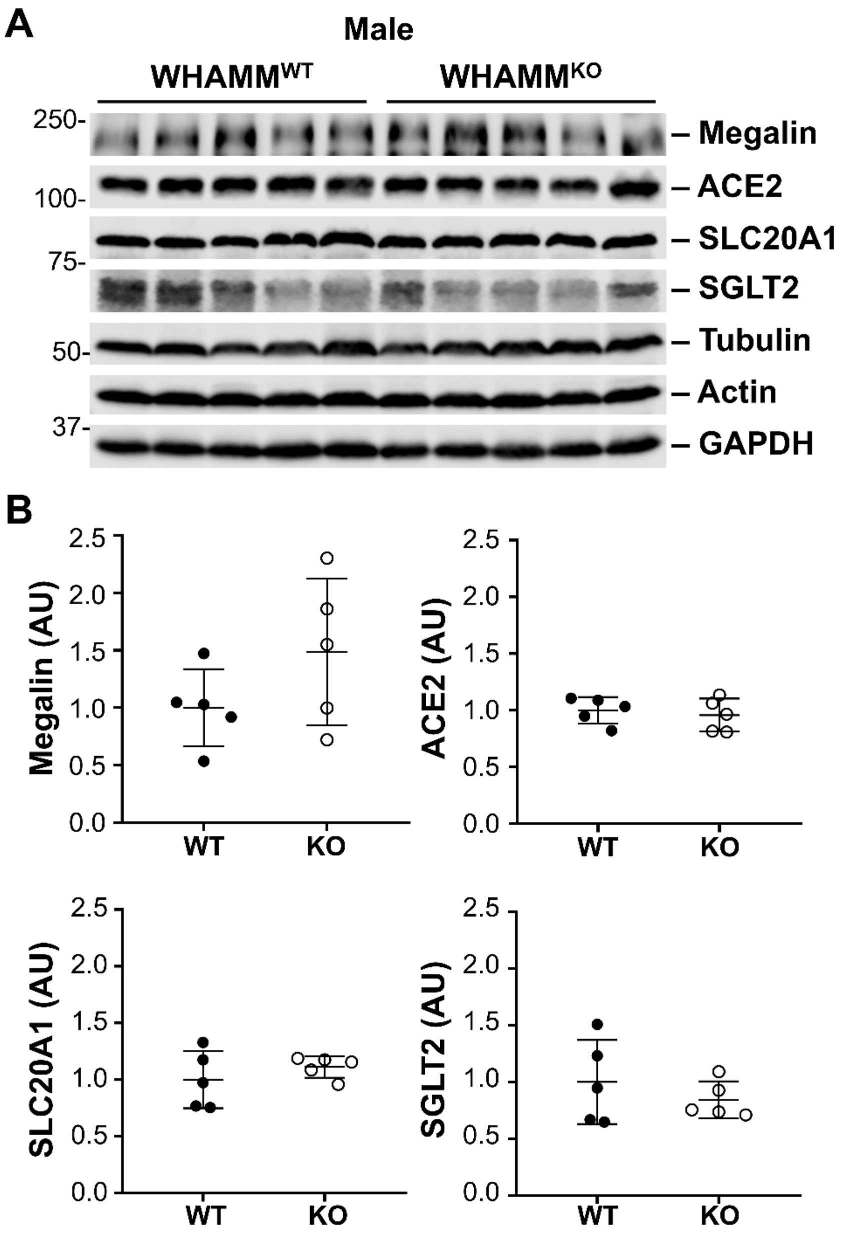

**Figure S4. Expression of several plasma membrane receptors is similar in male WHAMM$^{WT}$ and WHAMM$^{KO}$ kidneys. (A)** Kidney extracts from 5 male wild type (WHAMM$^{WT}$) and 5 WHAMM knockout (WHAMM$^{KO}$) mice were processed, and 50μg samples were subjected to SDS-PAGE and immunoblotted with antibodies to Megalin, ACE2, SLC20A1, SGLT2, tubulin, actin, and GAPDH. **(B)** Band intensities for the receptors in (A) were quantified relative to tubulin, actin, and GAPDH, and the normalized values were plotted. Each circle represents one mouse. Statistical bars represent the mean ±SD from n=5 mice.

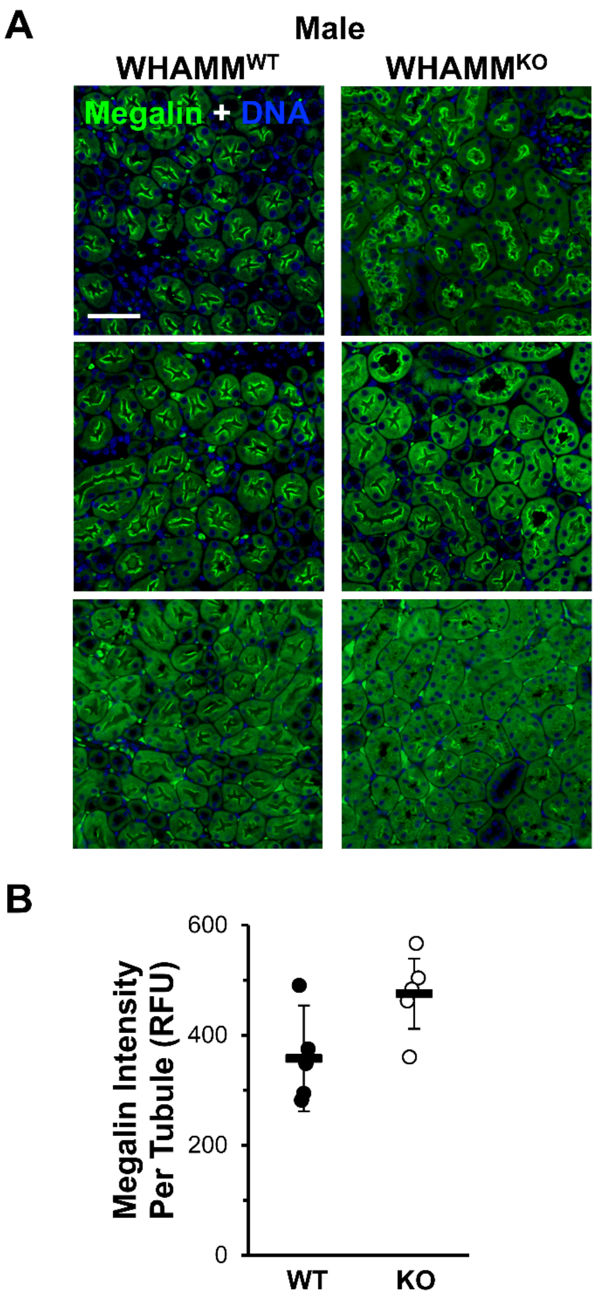

**Figure S5. Megalin localization and expression in kidneys from WHAMM$^{WT}$ and WHAMM$^{KO}$ mice. (A)** Kidney tissue sections from WHAMM$^{WT}$ or WHAMM$^{KO}$ male mice were stained with Megalin antibodies (green) and DAPI (DNA; blue). Scale bar, 50µm. **(B)** The Megalin fluorescence intensity per tubule was calculated in ImageJ. Each circle represents Megalin kidney staining from an individual mouse in which approximately 60 tubules were examined. Statistical bars represent the mean ±SD from n=5 kidneys.

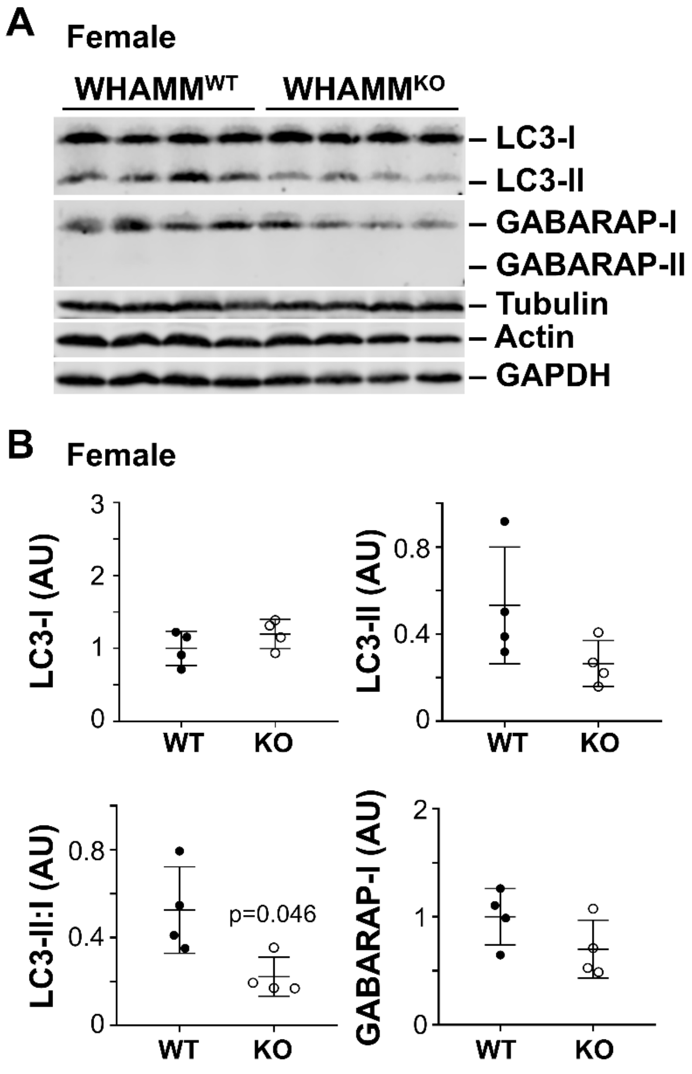

**Figure S6. The LC3 II:I ratio is slightly decreased in female WHAMM$^{KO}$ kidneys. (A)** Kidney extracts from 4 female WHAMM$^{WT}$ and 4 female WHAMM$^{KO}$ mice were processed, and 50µg samples were subjected to SDS-PAGE and immunoblotted with antibodies to LC3, GABARAP, tubulin, actin, and GAPDH. **(B)** LC3 and GABARAP band intensities were quantified relative to tubulin, actin, and GAPDH, and the mean normalized values were plotted. The LC3-II:I ratio was calculated by dividing the LC3-II band intensity by the LC3-I band intensity within each non-normalized sample. Each circle represents one mouse. Statistical bars represent the mean ±SD from n=5 mice. A significant p-value is noted (unpaired t-test).

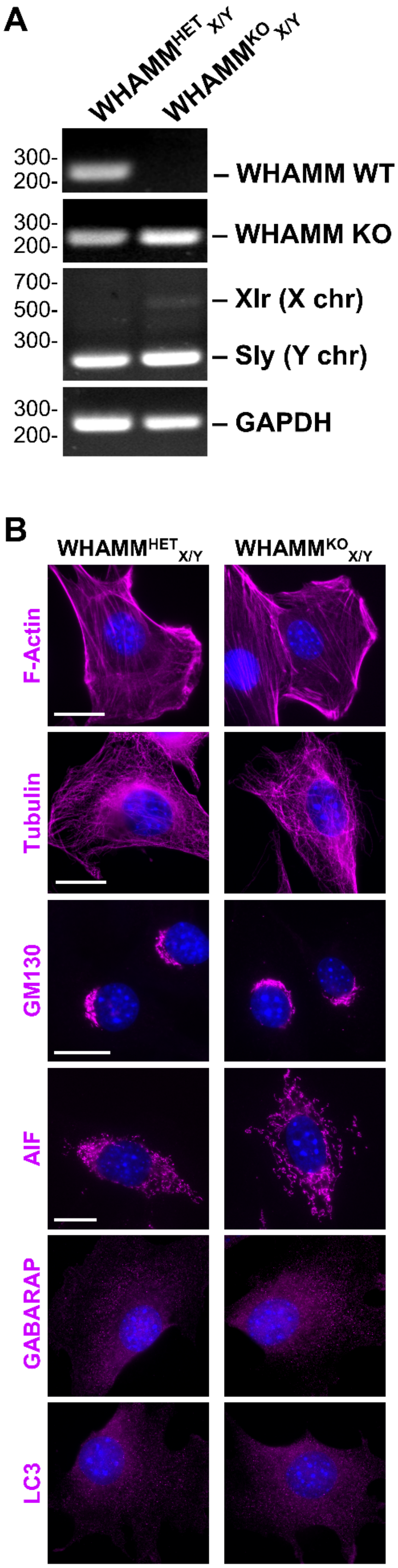

**Figure S7. Genotyping and microscopic analyses of mouse embryonic fibroblasts. (A)** Genomic DNA was isolated from mouse embryonic fibroblasts (MEFs) and subjected to PCR with primers to wild type *Whamm* (WHAMM WT), mutant *Whamm* (WHAMM KO), *Sly* (Y chromosome)/*Xlr* (X chromosome), and *Gapdh*, and visualized following agarose gel electrophoresis. **(B)** Male heterozygous ($WHAMM^{HET}_{X/Y}$) and male knockout ($WHAMM^{KO}_{X/Y}$) MEFs were fixed and stained with phalloidin to visualize F-actin or with antibodies to visualize microtubules (Tubulin), the *cis*-Golgi (GM130), mitochondria (AIF), or autophagosomes (LC3, GABARAP) each shown in magenta. DAPI-stained nuclei are shown in blue. Scale bar, 20μm. Despite multiple attempts, we were unable to isolate male wild type MEFs.

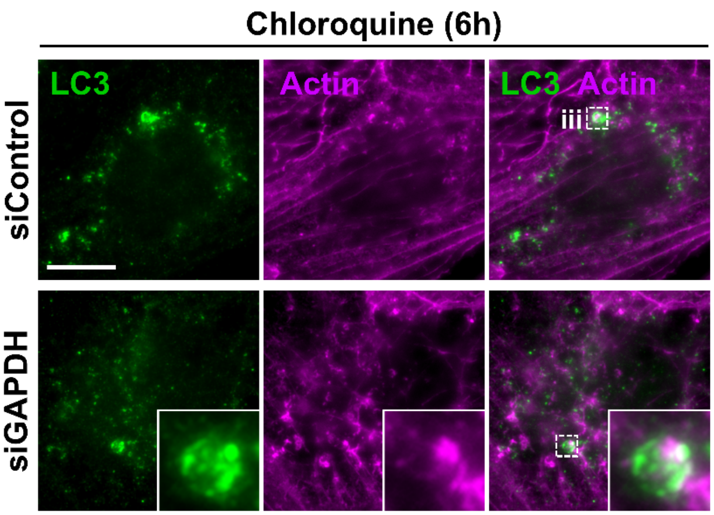

**Figure S8. Actin localizes to autophagosomes in proximal tubule cells.** HK-2 cells were treated with Control or GAPDH siRNAs and exposed to media containing chloroquine before being fixed and stained with antibodies to LC3 (green) and actin (magenta). Scale bar, 10μm. The magnification of (iii) is shown in Figure 3C.

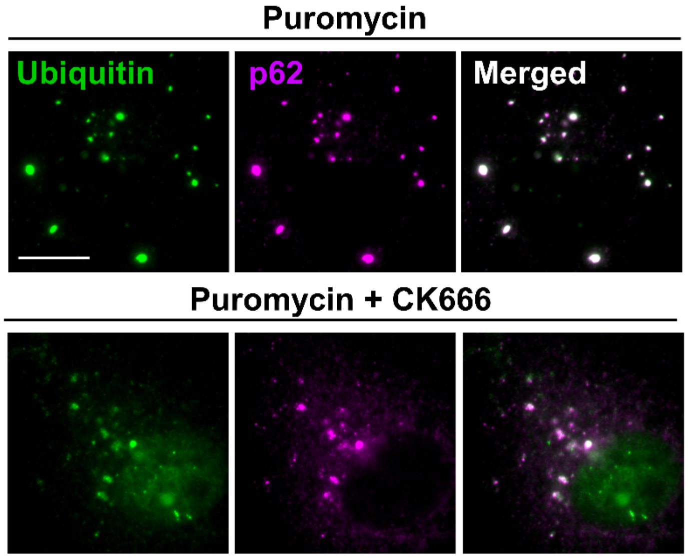

**Figure S9. Arp2/3 complex inhibition does not prevent the recruitment of p62 to ubiquitinated protein aggregates following proteotoxic stress.** HK-2 cells were treated with puromycin or puromycin plus CK666 for 2h before being fixed and stained with antibodies to ubiquitin (green) and p62 (magenta). Scale bar, 10µm. Exposure to puromycin increased the formation of bright, ubiquitin and p62-stained foci. Cells co-treated with puromycin and CK666 also harbored diffuse cytoplasmic ubiquitin and p62 staining, consistent with previous findings that the Arp2/3 complex is important for the formation of p62 bodies (Feng *et al.*, 2022). While the Arp2/3 complex may affect the coalescence of ubiquitin- and p62-bound cargo, it is not required for the recruitment of p62 to ubiquitinated material.
